# A composited genome approach to analyse cistrome diversity in barley

**DOI:** 10.64898/2026.10.27.762185

**Authors:** Amelie Kok, Jia-Ying Zhu, Natalie Bajaczyk, Vivien Joisten-Rosenthal, Sohini Mukherjee, Sergius Weizel, Venkata Suresh Bonthala, Nafiseh Sargheini, Bruno Huettel, Björn Usadel, Julia Engelhorn, Benjamin Stich, Thomas Hartwig

## Abstract

**Motivation:** Functional regulatory variants are important determinants of heritable diversity, yet their systematic identification requires both genome-wide maps of transcription factor (TF) occupancy and suitable genome sequences for mapping. While chromosome-scale assemblies remain costly for large association panels, short-read sequencing data are widely available. We investigated whether draft genomes reconstructed from short reads are sufficient for comparative MOA-seq analyses of TF occupancy.

**Results:** We generated genome-wide TF footprint maps for the barley cultivars Bowman, Morex, and Golden Promise, identifying approximately 240,000 loci per genotype. Roughly half of all footprints occurred in intergenic regions. Footprints conserved between genotypes were depleted for sequence variation and located closer to genes. Meanwhile, genotype-specific footprints were enriched for SNP-containing and gene-distal regions, highlighting the extensive diversity of regulatory landscapes among barley cultivars. We further compared two short-read-based draft genome assembly strategies: i) a SNP-replacement and ii) a Practical Haplotype Graph (PHG)-based composite genome approach, to enable MOA-seq analyses without chromosome-scale genome assemblies. Compared to the SNP-replacement strategy, the composite genome displayed 2.3-fold higher local sequence concordance for SNPs/INDELs relative to the published reference. Both draft genomes accurately reproduced MOA-seq profiles, recovering approximately 80 % of reference peak bases with per-base coverage correlations of 0.79–0.90 relative to the assembled reference. While the composite genome performed best across the genome overall, the simpler SNP-replacement strategy more accurately recovered TF occupancy within regulatory peaks due to fewer structural alignment artefacts.

**Availability:** Pipelines consist of existing open-source tools, Bash, GAWK and Java, and are available at [zenodo; tbd].

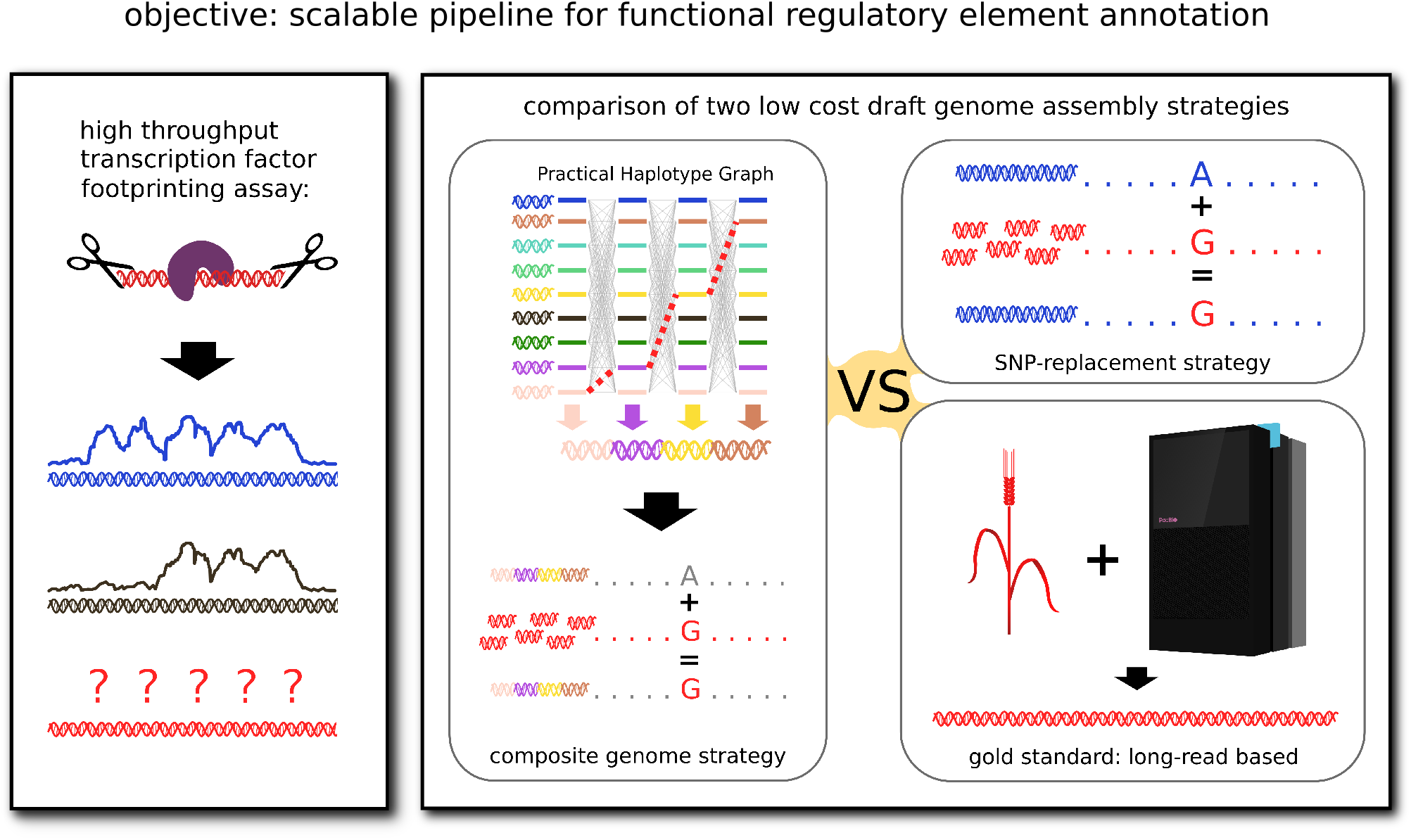

## Introduction

Genetic variation is the driving force of adaptation and the exploration of this variation is fundamental to plant and animal breeding (1). As such, methods to link genetic variation to phenotypic differences such as genome-wide association studies (GWAS) or quantitative trait locus (QTL) mapping (2–13), have become established and widespread. Previous studies suggested that about half of the genetic variation linked to phenotypic traits appears to be located in the non-coding space of the genome (14–17), which, to date, remains poorly annotated in most species, particularly when it comes to regulatory regions in plants (18, 19). An annotation of these regions could greatly accelerate the identification of the causal sequence variants of phenotypic traits, as has been shown in humans (20–22). To date, the barley regulatory space remains sparsely characterised compared to *Arabidopsis* or maize, despite its role as model organism for the cereals, with descriptions of broad accessible regions available for a few samples and genotypes (23, 24), but with larger-scale, comparative studies still pending.

Recently, a method called MNase-defined cistrome Occupancy Analysis (MOA-seq) has shown promise by providing a scalable, high-resolution approach for uncovering cistromes by locating putative transcription factor binding sites (TFBS). MOA-seq shares high overlap with TFBS determined through chromatin immunoprecipitation sequencing (ChIP-seq) (25, 26), the current gold-standard method for single-transcription-factor binding site localisation. Uncovered functional regulatory variants also co-located with genomic regions significantly associated with phenotypic variation through GWAS, and therefore have great potential to serve as targets for molecular breeding or genome editing (26). MOA-seq, like other approaches, relies on DNA-sequence mapping. As such, its accuracy depends on the quality of the underlying reference sequence (26), to an as of yet unknown degree. This aspect gains even more importance when trying to quantify differences between samples or genotypes.

While a genome assembly using long-read sequencing has become more affordable, with an increasing number of pan-genomes available (27–32), it can still be prohibitively expensive for projects that involve hundreds or thousands of genotypes, which are common for projects in the field of quantitative genetics. Therefore, an interest exists in developing cost-effective strategies to assemble draft-quality genomes, which may provide sufficient quality when used in a sequencing-based assay, e.g. for the annotation of the non-coding space of the genome such as with MOA-seq or ChIP-seq. While the use of one shared, potentially modified, sequence is feasible, results may be skewed by reference bias. This issue could be alleviated by creating personalised genome sequences through integration of multiple existing sequences.

Pan-genome graphs can capture the combined sequence information of populations in a single data structure (33–36). They can be used to compare genomic variation across assemblies and potentially to construct a personalised composite genome. Commonly used, so-called variation graphs, e.g. minigraph-cactus, PGGB, PanGenie (33–35), where each variant creates a new node in the graph structure, are currently computationally feasible only for smaller genomes. Jayakodi et al. (27) demonstrated that a minigraph-cactus-based graph is possible for barley, with a medium-sized genome of ~4.2 Gbp, by limiting it to individual chromosomes and variants greater than 50 bp. This indicates that applicability likely breaks down for larger genomes. The Practical Haplotype Graph (PHG) (36) is a light-weight pan-genome graph with a simplified structure, designed for use with larger and complex plant genomes. Designed for genotyping, it sacrifices some accuracy for performance by having nodes consisting of longer sequences (haplotypes) corresponding to a predefined set of genomic ranges in a reference genome (reference ranges).

In this study, we aim to explore different strategies for performing large-scale cis-element discovery through MOA-seq in the future, as well as provide a first basic description of cistrome variation in barley. We constructed a PHG-based on available barley genome assemblies (37), used its pathfinding algorithm to extract a set of haplotypes closely related to a genome of interest (Bowman), and built a personalised composite assembly. We compared this composite genome with a published assembly of the Bowman cultivar based on long-read and Hi-C sequencing and with a SNP-replaced genome based on a single assembly. We characterised sequence variants between the three sequences by type and potential origin and evaluated the performance of both the composite genome and the SNP-replaced genome regarding their abilities to facilitate an accurate analysis of MOA-seq data. Additionally, we provided genome-wide sets of putative TFBS for three important barley genetic stock, namely Morex, Bowman and Golden Promise, to exemplify downstream cistrome diversity analyses.

## Results

### TFBS discovery through MOA-seq in three barley genotypes

In order to characterise genome-wide transcription factor (TF) footprints in barley, we performed MOA-seq for three important genetic stocks of barley (*Hordeum vulgare*): Bowman, Morex and Golden Promise. MOA-seq was performed on leaf tissue for three biological replicates per inbred. The resulting read coverage correlated well among replicates, with all exceeding the ENCODE recommendation of a Spearman correlation coefficient above 0.8 (average: 0.94; lowest correlation between two Golden Promise replicates: 0.87; Supplementary Fig. 1; (38)). After merging read alignments of the replicates for an increased sensitivity when calling peaks, we observed an average of 243,082 putative TF binding peaks per inbred (compare Table 1) ranging between 232,618 and 253,709 for Morex and Bowman, respectively. About half (51.4 %) of these peaks were located in intergenic regions (i.e. *>*5 kb away from the closest gene). Promoter and terminator regions, up to 5 kb adjacent to transcription start and termination sites (TSS/TTS), contained about 34.2 % of MOA-seq peaks (Fig. 1 A). Untranslated regions (UTRs) harboured another 7.6 % of MOA footprints, while intron and exon overlaps both occurred at lower frequencies of 4.4 % and 2.5 %, respectively. For the 5 kb regions adjacent to the TSS and TTS as well as the UTRs, more MOA-seq footprints were located upstream than downstream of the coding region. For peaks located outside of genes, the median distance to the nearest gene ranged between 15 - 25 kb depending on the inbred line, indicating many long-range interactions (compare Fig. 1 B).

**Table 1.** Comparison of peak-calling in three barley inbred lines. Peak coordinates for each inbred were translated to each of the other coordinate systems (coord. syst.) to determine which peaks were shared among inbreds based on peak-calling. Overlaps were determined in each coordinate system individually for each comparison.

| peaks from | translated to | peaks called | peak-middles translated | overlapping in both coord. syst. |
| --- | --- | --- | --- | --- |
| Bowman | Golden Promise | 253,709 | 232,525 | 59,562 |
| Golden Promise | Bowman | 242,920 | 224,470 |  |
| Bowman | Morex | 253,709 | 231,855 | 85,543 |
| Morex | Bowman | 232,618 | 217,700 |  |
| Golden Promise | Morex | 242,920 | 224,113 | 58,091 |
| Morex | Golden Promise | 232,618 | 214,686 |  |

**Fig. 1.**
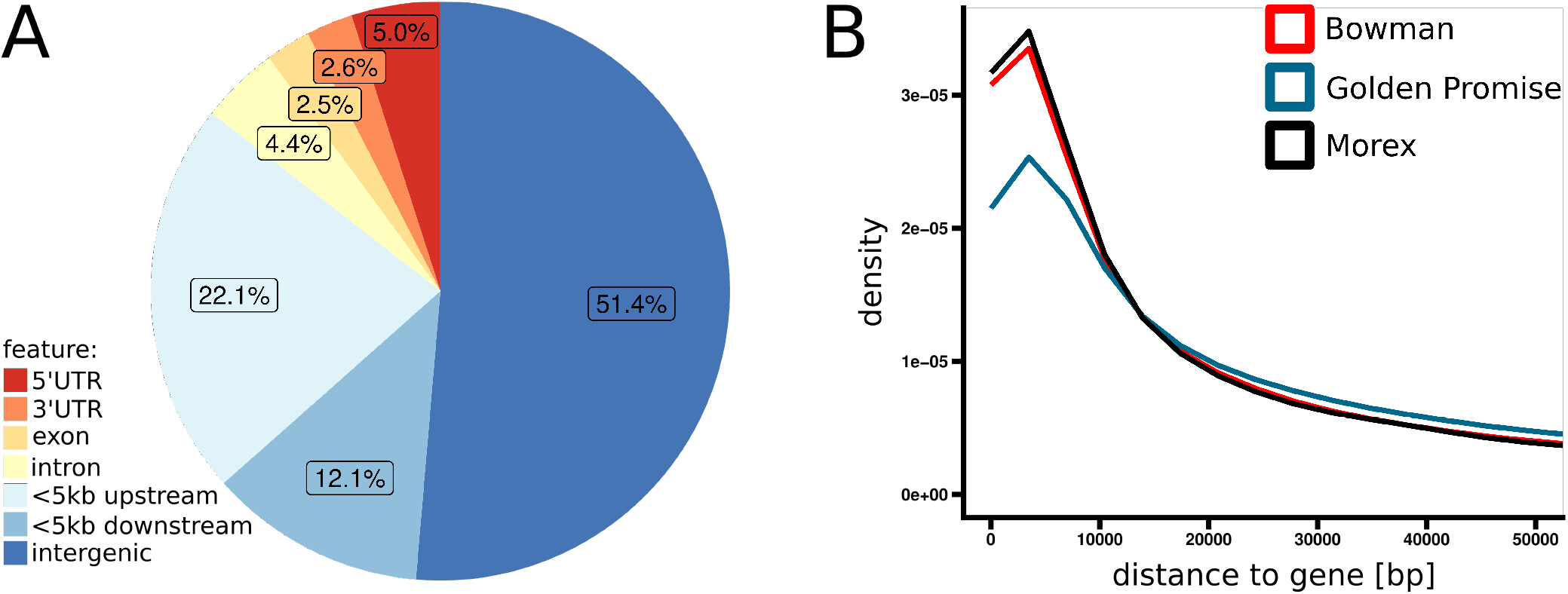
Localisation of MOA-seq peaks relative to gene annotations. **A** Number of peaks overlapping with the different genomic features relative to gene annotations. Plotted is the average between the three genotypes Bowman, Morex and Golden Promise. **B** Relative distance of all peaks to the nearest gene. Excluded are peaks overlapping with genic regions. The x-axis is truncated at 50 kb.

To investigate the conservation of peak distributions between genotypes, the positions of peaks were compared in a pairwise manner. Across all peaks, an average of 7.7 % could not be translated between the coordinate systems of two genomes, i.e. the peak centre coordinate was not alignable (Bowman vs. Golden Promise and the reciprocal comparison: 21,184/253,709 and 18,450/242,920, respectively; Bowman vs. Morex and the reciprocal comparison: 21,854/253,709 and 14,918/232,618, respectively; Golden Promise vs. Morex and the reciprocal comparison: 18,807/242,920 and 17,932/232,618, respectively), potentially representing presence/absence variation. The remaining peaks were conservatively classified as shared if a peak center overlapped a peak in the comparison genome, and vice versa (for details, see Methods). On average, 30.2 % of the remaining MOA peaks were found to be shared between genomes (Bowman vs. Golden Promise and the reciprocal comparison: 59,562/232,525 and 59,562/224,470, respectively; Bowman vs. Morex and the reciprocal comparison: 85,543/231,855 and 85,543/217,700, respectively; Golden Promise vs. Morex and the reciprocal comparison: 58,091/224,113 and 58,091/214,686, respectively) (Table 1).

To correlate shared versus non-shared peaks with DNA sequence variation and gene positions, we performed a detailed analysis on the translatable Morex and Bowman peaks. We found a significant depletion of shared peaks within peaks overlapping with variants, while non-shared peaks were enriched (Supplementary Table 1; P-value *<* 0.05 hypergeometric enrichment test). Shared peaks overlapping with SNPs further had a lower median count of SNPs per peak than peaks not categorised as shared (Fig. 2 A). Finally, shared peaks were also located closer to genes than peaks not categorised as shared (Fig. 2 B).

**Fig. 2.**
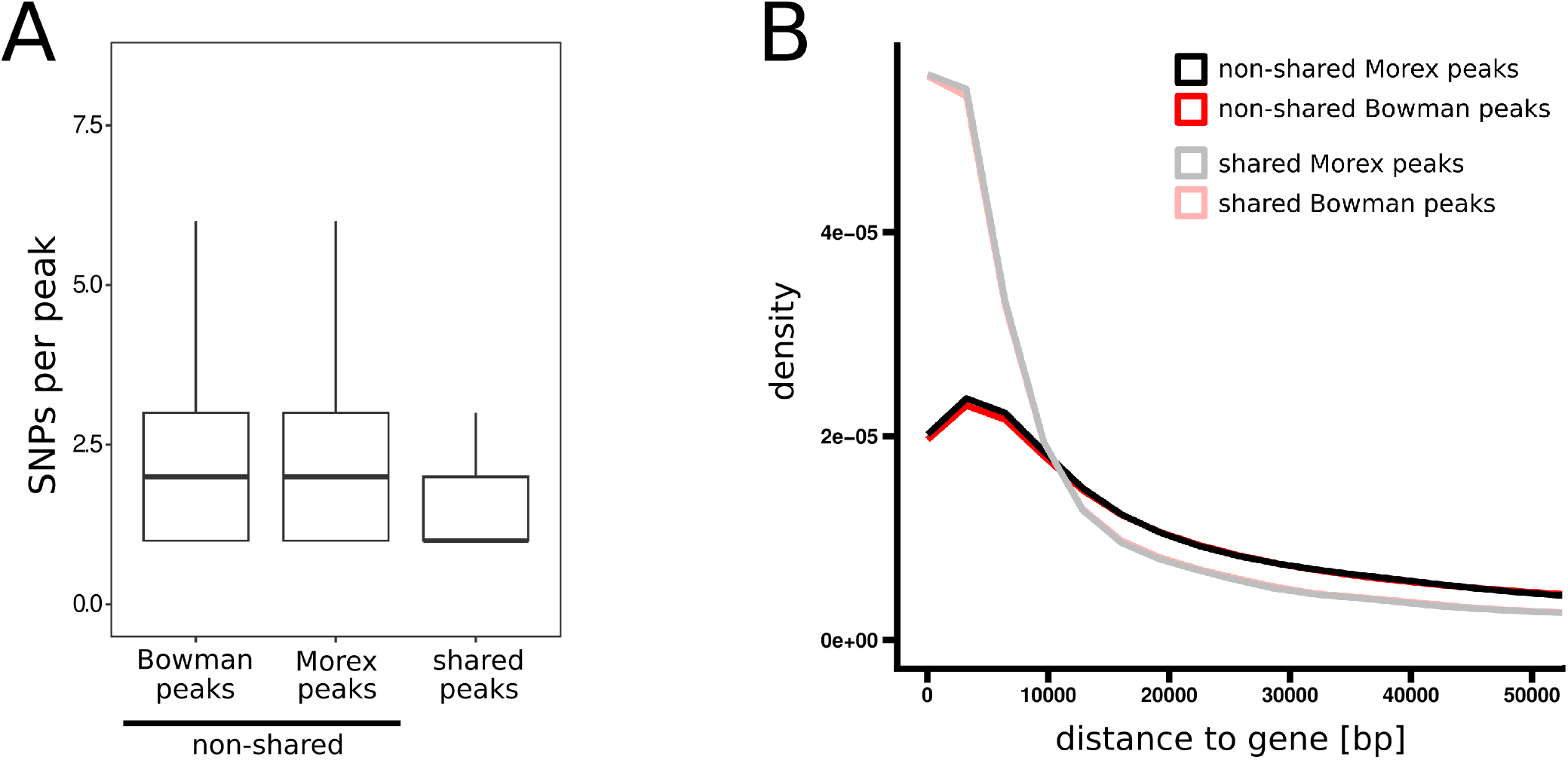
Characterisation of MOA-seq peaks categorised based on whether or not they were shared between Bowman and Morex. **A** Number of SNPs overlapping a peak. **B** Relative distance of peaks to the nearest gene. Excluded are peaks overlapping with genic regions and peaks are categorised as described above. The x-axis is truncated at 50 kb.

Larger variants, especially large insertions (*>*50 bp), could introduce TFBS, however, only 0.45 % of bases within large insertions were annotated as peaks, compared with 0.89 % across the remainder of the genome, indicating a depletion of TFBS within large insertions.

We not only observed shared and non-shared peaks on a genome-wide scale, but also on the level of individual, previously studied genes. For example, the promoter of *short rachilla hair 1* (*HvSRH1*; HORVU.MOREX.PROJ.5HG00435480.1, HORVU.BOWMAN.PROJ.5HG00322140.1, HORVU.GOLDEN_PROMISE.PROJ.5HG00440000.1; Fig. 3 A), a gene located within a GWAS interval linked to the rachilla hair phenotype (27, 37, 39), contained MOA-seq peaks shared among the three genotypes. Although the three genotypes differ in rachilla hair phenotype, this observation of a shared gene-proximal regulatory pattern matches the evidence and hypothesis of Jayakodi et al. (27) that differences in *HvSRH1* expression are caused by a distal regulatory element.

**Fig. 3.**
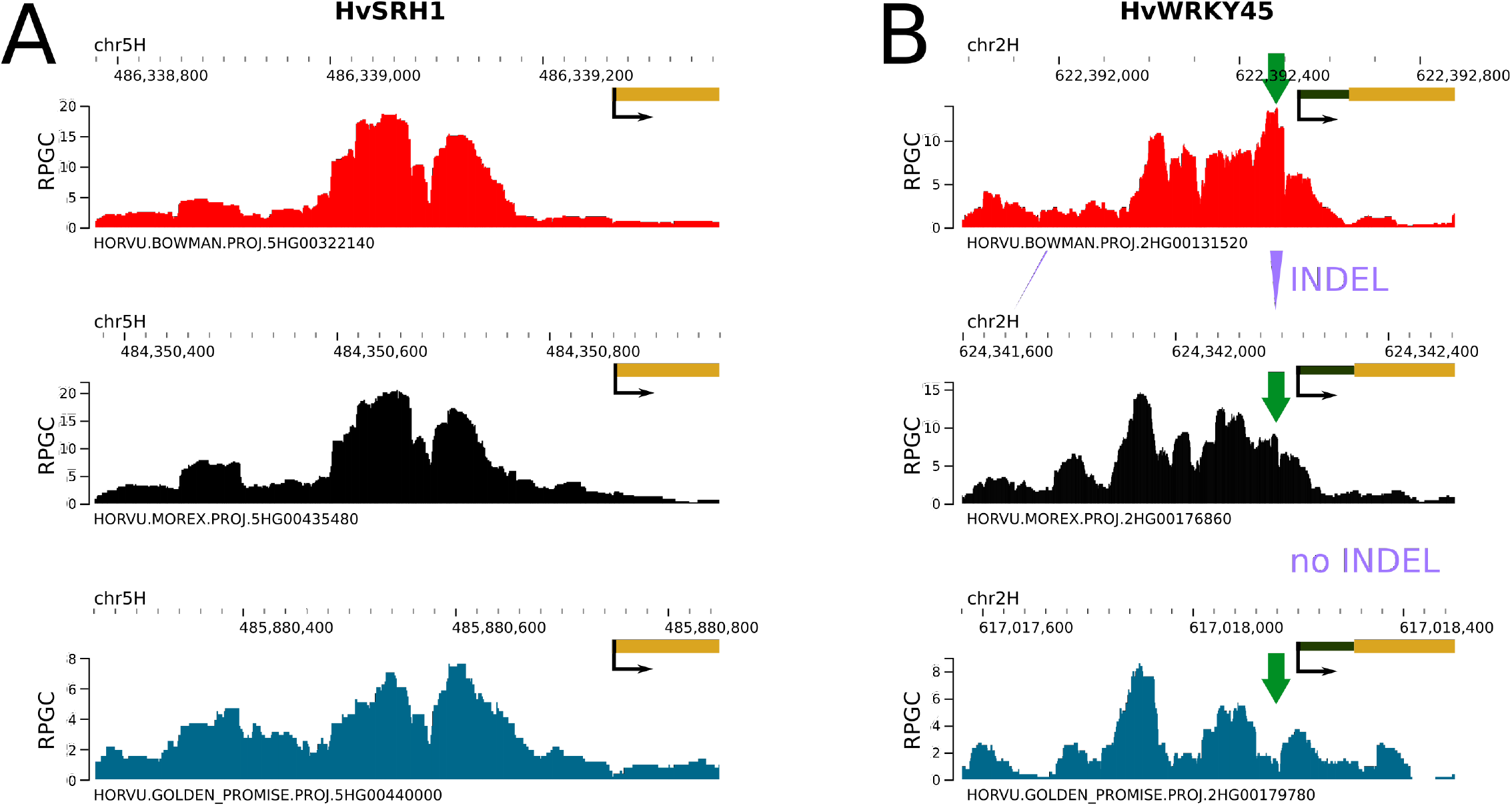
Two example peaks demonstrating similar and differential signal among three inbreds. Browser view of MOA-seq peaks near *HvSRH1* (**A**) and *HvWRKY45* (**B**) genes. Raw read coverages were normalised using RPGC (reads per genomic content) methods. Purple triangles represent INDELs. Green arrows indicate a change in MOA-seq signal at the INDEL.

In contrast, MOA-seq peaks within the promoter of *HvWRKY45* displayed a difference in signal strength overlapping with a 29-bp insertion (Fig. 3 B). This insertion contains an ‘AGCCACC’ Box S *cis*-regulatory element and its presence has previously been found to correlate with the differential expression of the adjacent *HvWRKY45* gene in two wild barley populations (40). Here, we found that Bowman shares the 29-bp insertion allele with the wild barley populations, and observed a spike in MOA-seq signal in Bowman compared to the other two genotypes. A comparative gene expression or mutational assay would, however, be required to prove a direct link between the sequence variant, TF occupancy and gene expression. Overall, these two examples provide a small insight into the varied landscape of TF binding in barley and exemplify the potential of MOA-seq in pinpointing functional variation in cis-regulatory regions.

### Assembly of draft genome sequences from short reads for population-scale TFBS comparisons

Effective study of functional variation in *cis*-regulatory regions not only requires scalable approaches for genome-wide characterisation of TF footprints like MOA-seq, but also methods for genome assembly of sufficient accuracy. Especially for plants with medium to large genome sizes like barley, long-read sequencing data for a large number of inbred lines remain a significant cost factor while short-read sequencing data are often available from genotyping projects. To facilitate MOA-seq analysis in large-scale projects, we aimed to compare two approaches of differing computational complexity to establish draft genome sequences from readily available short-read sequencing data. To validate the approaches, we used short-read sequencing data of the Bowman genotype, for which an assembled genome is available, for comparison. As a first approach for draft genome assembly, we tested a SNP-replaced genome sequence by integrating called SNPs into the genomic sequence of the *Hordeum vulgare* reference cultivar Morex (41). The SNPs for this approach were called using 30x coverage whole genome shotgun sequencing (WGS) (42) and the Genome Analysis ToolKit (GATK; (43)).

For the second approach, termed composite genome approach, we tested the iterative construction of a genome sequence based on both SNPs and other genomic variants. To do so, 19 publicly available genomes (37) were individually aligned against the Morex genome (v3; (41)) and a Practical Haplotype Graph (PHG, (36)) was constructed from the alignments (compare Fig. 4 step 1). Haplotypes for genome construction were then selected using the PHG software’s pathfinding based on an alignment of the Bowman WGS reads to the graph (compare Fig. 4 step 2). To test parameter selection (see Methods) and the ability of the pathfinding algorithm to identify haplotypes with the highest similarity to the input WGS reads, pathfinding was first performed based on ten different, previously published sets of WGS reads belonging to various barley cultivars (44). Overall, the frequency with which haplotypes of the 19 cultivars were selected in the path, broadly correlated with the sequence dissimilarity of each of the 19 cultivars compared to each of the cultivars of the ten published datasets (measured as modified Rogers distance; −0.79 Spearman correlation coefficient; see Fig. 5 A). As expected, this illustrates that more closely related haplotypes were selected more frequently than those of more distantly related cultivars. This indicated the ability of the pipeline to select haplotypes that are similar to the input WGS reads’ sequences. After performing the pathfinding based on Bowman WGS short-reads, the selected haplotypes were concatenated to form chromosome-level preliminary sequences in the order of the Morex guide haplotypes. To further improve local sequence context, we integrated both SNPs and INDELs (*<*50 bp), called based on the same Bowman WGS reads used to generate the SNP-replaced genome, into the preliminary composite sequence (compare Fig. 4 step 3). We refer to the resulting sequence as “composite genome” and the preliminary sequence as the “preliminary composite” sequence.

**Fig. 4.**
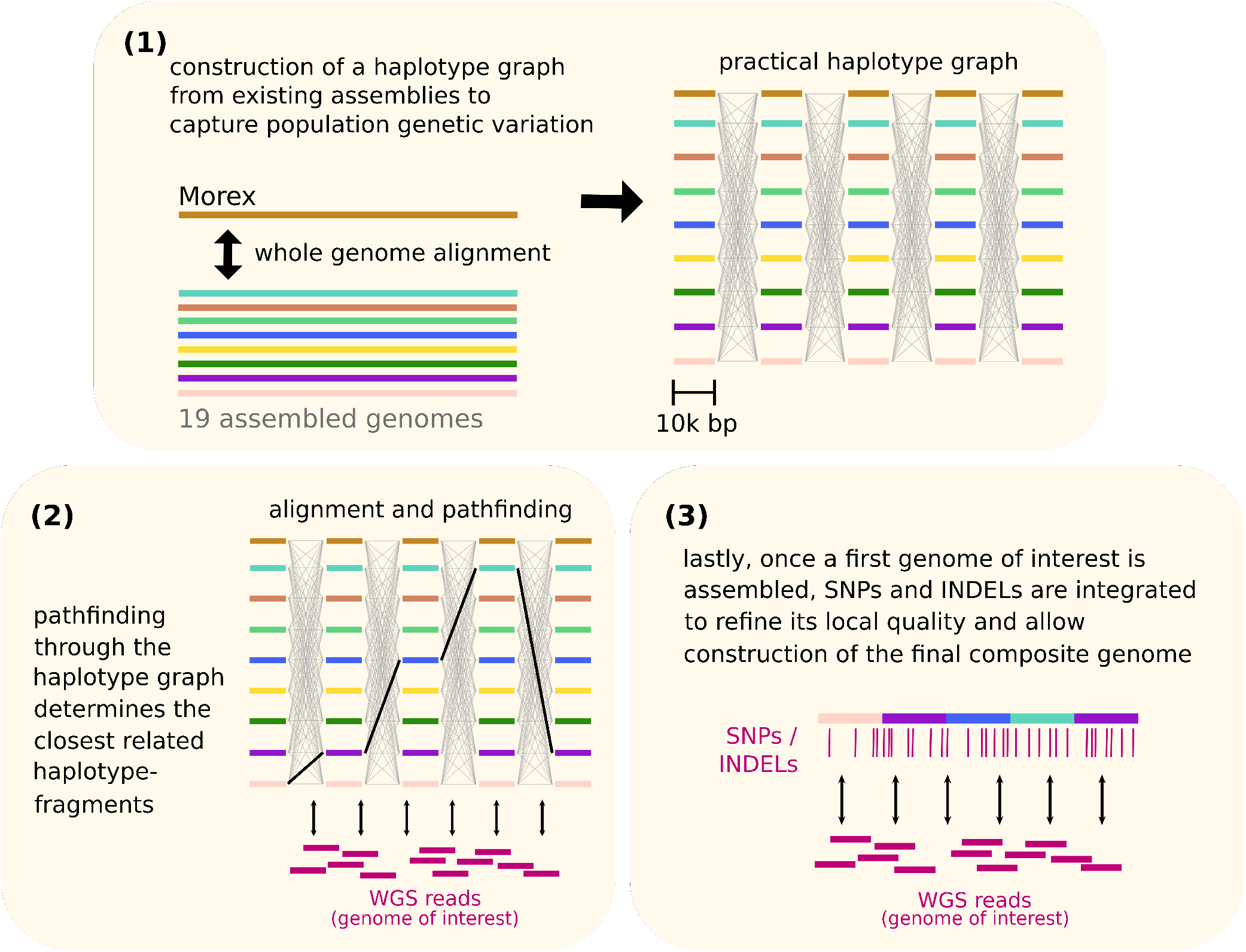
Schematic overview explaining composite genome creation for a genome of interest. Step 1: A practical haplotype graph (PHG; (36)) was constructed from pairwise whole-genome alignments of 19 publicly available genomes (37) against the Morex v3 genome (41). A reference range length of 10 kb each was selected as the basis for computing graph nodes. **Step 2**: Short whole-genome sequencing reads of a genome of interest (here: Bowman) were aligned against the graph and the optimal path was determined through the pathfinding algorithm. **Step 3**: Haplotypes selected through pathfinding were concatenated into the preliminary composite sequence, which was then used as a base for variant calling using GATK. SNPs and INDELs that were called were then integrated into the preliminary sequence to form the finalised composite genome sequence.

**Fig. 5.**
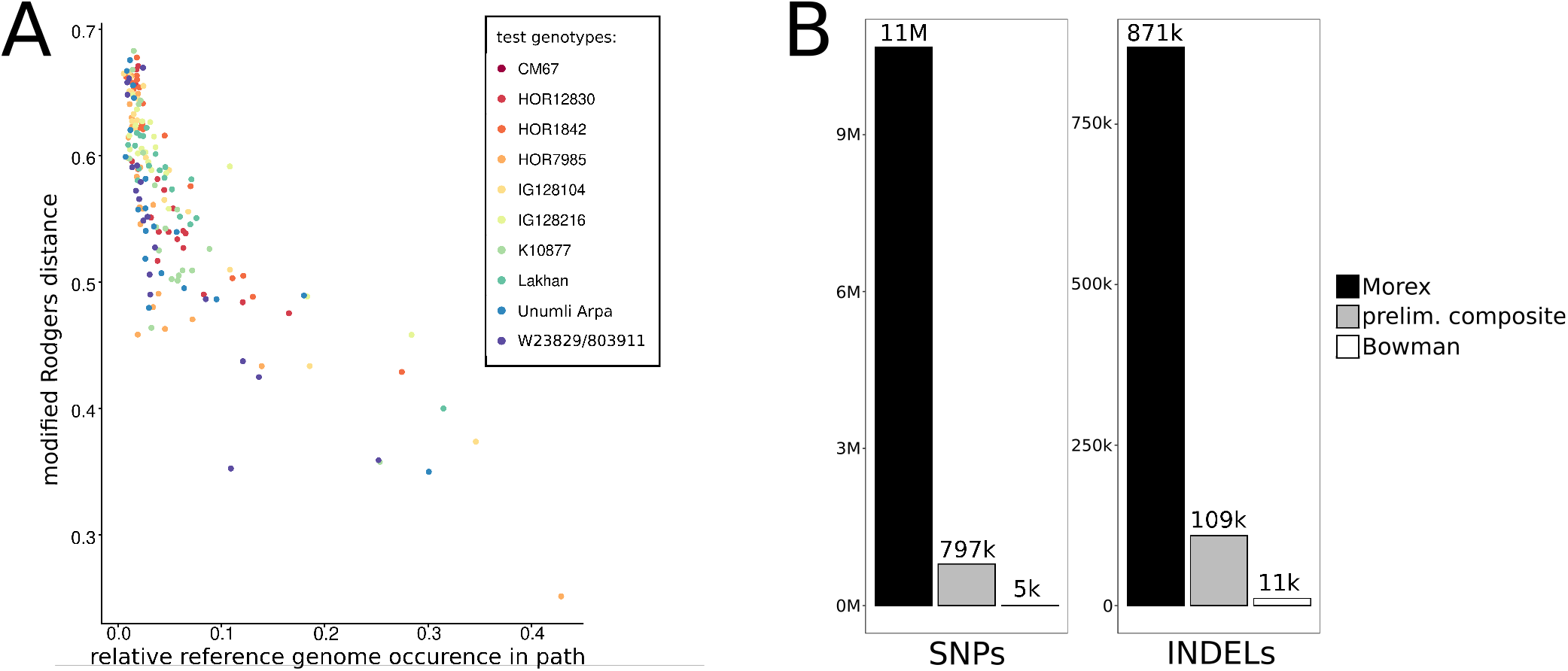
Intermediate evaluation of the composite genome creation pipeline. **A** Performance test of the PHG pathfinding algorithm using short whole-genome sequencing reads of ten barley genotypes of the parental inbreds of the double round-robin populations (45). For each of the ten test inbreds, the modified Rogers distance was determined relative to each of the 19 genomes that made up the PHG. Additionally, the frequency of how often a haplotype of each of the 19 genomes was selected through pathfinding was determined for each of the ten tests. **B** Number of SNPs and INDELs called using GATK and Bowman short-reads against the Morex genome, preliminary composite genome and Bowman reference genome.

### Sequence-level comparison of draft genome assemblies

We hypothesised that the preliminary reference sequence more closely resembled that of the genome of interest, thereby providing a more suitable mapping template for subsequent variant calling and integration. To evaluate this, we first compared the number of variants called with Bowman WGS reads against either the PHG-based preliminary composite or the Morex reference. We detected 14- and 8-fold fewer SNPs and INDELs in the preliminary composite sequence compared to the Morex sequence, respectively (Fig. 5 B). Overall mappability was higher and the base mismatch rate was lower for the PHG-based preliminary composite genome than for the Morex genome sequence, respectively (Supplementary Table 2). Together, these observations indicate that the composite genome strategy indeed generates a preliminary sequence closely related to the genome of interest.

We then compared the final composite sequence after variant integration to the SNP-replacement strategy, where SNPs were directly integrated into the Morex reference, regarding their ability to correctly capture SNPs. When using a truth set of about 19 M SNPs generated through whole-genome alignment (WGA) between the Bowman and Morex genomes, the SNP-replaced genome recovered about 53 % (~10 M) of this truth SNP-set. The composite genome sequence, on the other hand, correctly captured about 79 % (~15 M) of the truth SNP-set, displaying a higher sequence similarity to the Bowman reference with regard to SNPs (Fig. 6). These results indicate that the composite genome strategy can reduce this limitation by creating a more closely related intermediate template first.

**Fig. 6.**
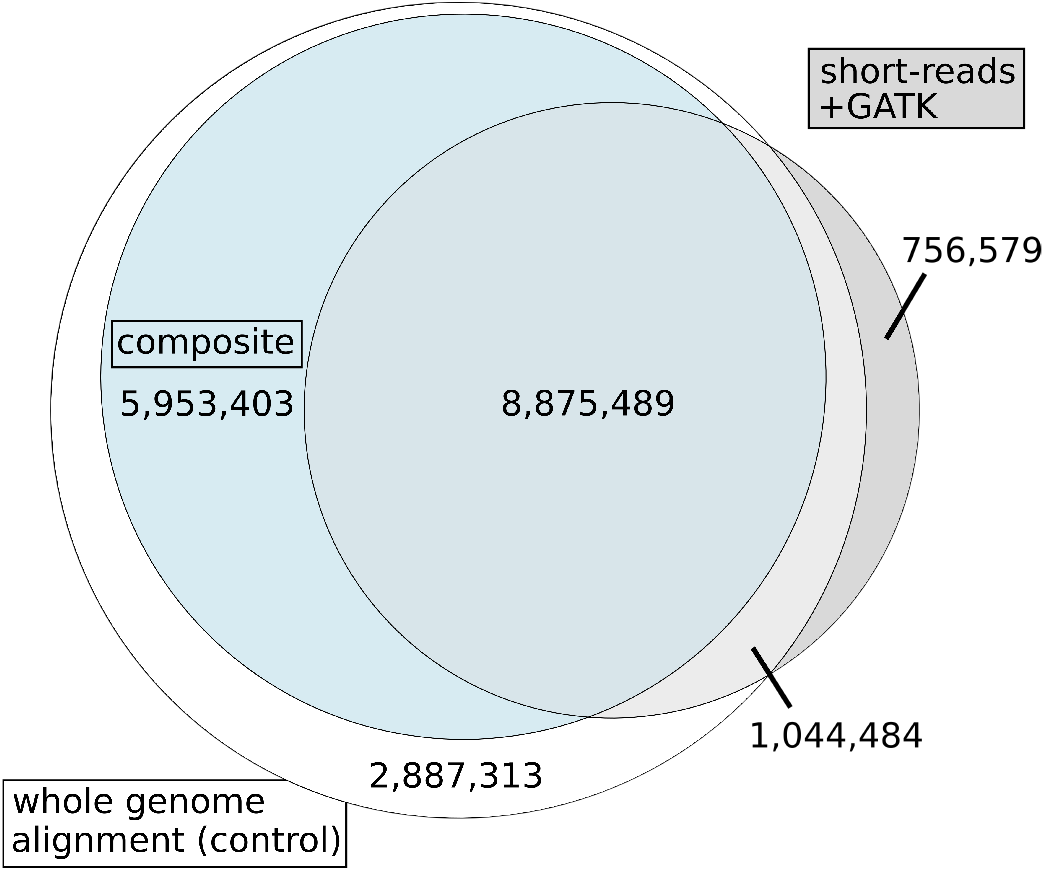
Ability of short-read-based variant calling to capture SNPs. Number of SNPs called between the Bowman and Morex published reference genomes. White circle: number of SNPs called between Bowman and Morex using whole-genome alignment. Grey circle: number of SNPs called between Bowman and Morex using short-reads of Bowman and GATK after aligning reads to Morex. Blue circle: subset of the white circle where alleles were matching between the Bowman reference and the composite genome sequence. Overlaps show SNPs determined by two or all three methods.

In a second comparison, the sequences of both draft genomes after variant integration were directly compared to the Bowman reference using WGA. Variants were called from pairwise alignments of each draft genome to the reference sequence (Table 2). The majority of bases, about 3.8 Gbp or ~90 % of the Bowman reference, were determined to be non-variant in both WGAs. Also shared between both WGAs, the majority of variant basepairs were part of structural variants (SVs), likely due to their size. On the other hand, we also observed differences between the

**Table 2.**
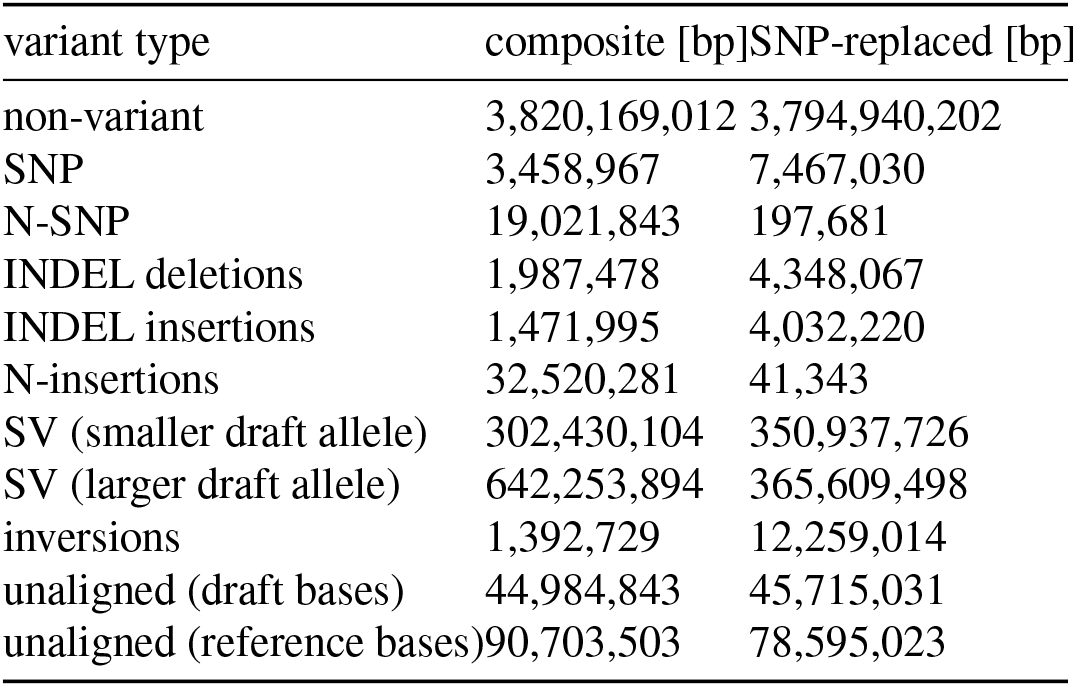
Variant calling between the two draft genome sequences and the Bowman reference sequence. Cumulative length of each variant type. In case of types with differing lengths for each allele (INDELs, insertions, deletions, duplications), the length of the longer allele was added to the sum. INDEL insertions and deletions, as well as N-insertions were determined for the draft allele, i.e. the draft allele displaying the given variant type. N-SNPs are single base pair changes to the ambiguous base “N”. N-insertions are insertions where all inserted bases were called as “N”. Under inversions, only inversions without internal smaller variants are listed. For unaligned regions bases unaligned in either genome were counted.

WGAs. The WGA of the composite genome showed: i) a notable increase of non-descriptive bases (“Ns”), ii) more unalignable regions, and iii) a larger number of bases within SVs called against Bowman, compared to the SNP-replaced genome. These observations were not unexpected: To reduce computational workload, the thousands of unplaced scaffolds for each PHG genome were connected via 1000 Ns, in addition to the relatively large number of Ns included in the PHG genomes compared to the highly resolved third iteration of the Morex genome with far fewer Ns. Choosing relatively small PHG reference ranges of 10 kbp length meant that they were spread across less conserved non-coding regions. Furthermore, larger SVs between the PHG genomes and Bowman, which span multiple ranges, would be difficult to place correctly, potentially resulting in multiple SVs. These factors made it more challenging to resolve large SVs and transitions across ranges. On the other hand, the WGA comparison again showed that the composite sequence was more similar to Bowman than the SNP-replaced sequence in local sequence contexts. In particular, we identified fewer SNPs (3.5 M vs. 7.5 M) and INDELs (3.5 M vs. 8.4 M bp) for the composite than the SNP-replaced genome compared to the assembled Bowman reference. In addition, the positions of the SNPs and INDELs were mostly unique to each WGA, with about 195 k SNPs and 157 k INDELs having shared positions and *>*1.8 M and *>*3.5 M INDELs and *>*2.5 M and *>*5.3 M SNPs having unique positions (Supplementary Table 3), indicating that short variants likely originate from different sources in each draft genome. Taken together, these comparisons indicate that, while the composite genome showed a higher sequence similarity to the Bowman reference sequence in local contexts (SNPs/INDELs) than the SNP-replaced genome, it also appeared to include more sections that are difficult to align or are larger structural variants.

To determine the origin of regions that were unalignable in the composite WGA but alignable in the SNP-replaced WGA, additional analyses were performed. These un-alignable regions formed 27 continuous segments, with approximately 30 % of their base pairs located within 40 Mbp of the chromosome ends (Supplementary Table 4), even though these terminal regions account for only about 13 % of the genome. One possible explanation is that, for some reference ranges used by the PHG to partition the Morex genome, the pathfinding algorithm failed to select a haplotype. However, only about 13 % of these bases unalignable in the composite WGA but alignable in the SNP-replaced WGA were located within reference ranges for which no haplotype had been selected.

Another property of the composite genome sequence was the large count of the non-descript base “N” mentioned above. To investigate the cause, the number of “N” for each of the haplotypes selected by the pathfinding algorithm was determined (Supplementary Table 5). While the majority (72 %) of the approximately 203 million “N” bases in the preliminary composite sequence stemmed from the chromosomal regions of the original assemblies, almost a third of the “N” originated from the scaffolds, which were concatenated with 1k “N” as a divider to reduce computational complexity in the composite genome pipeline. This occured despite scaffold sequences only being selected by the pathfinding in about 300 cases. The final step of the pipeline, i.e. the integration of variants into the preliminary composite sequence, reduced the number of “N” from about 203 million down to about 147 million. Thus, both the large number of “N” and the unalignable regions are remaining obstacles to the creation of an accurate genome sequence using this composite genome strategy.

### Short-read-based assembly strategies recover around 80 % of bases called in MOA-seq peaks

We also wanted to understand if the composite or SNP-replaced genomes were more suitable for cistrome analysis compared to a single related reference genome. Both draft genomes were evaluated as templates for transcription factor foot-printing analysis, specifically for accurately reproducing MOA-seq read mapping to the Bowman reference sequence. Performing read mapping and peak calling against the two draft genome sequences showed that about 83.2 and 79.2 % of the base pairs called as peaks in the reference genome were recovered by the SNP-replaced and composite genome’s alignment and peak calling, respectively (Table 3). This matches the observation that the majority of peak bases were located within regions that were non-variant between the two draft genomes and the reference (Supplementary Table 3). About 20.7 and 20.6 % of the bases annotated as peaks in the SNP-replaced and composite genomes, respectively, were not called in the Bowman reference. However, peak calling thresholds were set per sample, and results therefore may not always represent a comparison of significant coverage differences. Instead, the results represent an evaluation of the two draft genomes for common genomic applications, such as peak calling. Together, our observations indicate that both draft genomes allowed a high recovery of peaks, but also a considerable rate of discordant peak calls.

**Table 3.** Peaks recovered using draft genomes. Statistics for peaks called in both draft genomes and the Bowman reference genome, as well as the percentage of bases that are called as peaks correctly (true positives) or incorrectly (false positives) in either draft genome when compared to bases called as peaks in the reference.

| genome | peak number | cumulative peak length [bp] | median peak width [bp] | true positives [%] | false positives [%] |
| --- | --- | --- | --- | --- | --- |
| Bowman reference | 280,701 | 53,266,757 | 126 | - | - |
| SNP-replaced | 279,093 | 55,878,209 | 133 | 83.17 | 20.72 |
| composite | 266,891 | 53,088,638 | 132 | 79.18 | 20.56 |

For a more direct comparison of each genome’s ability to facilitate accurate MOA-seq analysis, we determined quantitative differences between the MOA-seq coverage in the Bowman reference and the two draft genomes. To this end, the read coverage per base pair of the reference sequence alignment was compared to the coverage of that base pair in the alignments against the two draft genome sequences (limited to chr1H to save computational resources). The coverage of both draft genome sequences correlated well with that of the reference sequence, though the composite genome performed slightly better, with a correlation coefficient of 0.84, compared to the SNP-replaced genome with a correlation coefficient of 0.82 (Spearman’s rho; Table 4). Both draft genomes performed better than using the Morex reference genome for read alignment, which yielded a correlation of 0.76. When limiting the investigated bases to only bases in Bowman reference peaks or peak summits, however, the SNP-replaced genome performed best with correlation coefficients of 0.90 and 0.88 for peak bases and peak summit bases compared to 0.88 and 0.86 for the Morex sequence and 0.80 and 0.79 for the composite (Table 4; Fig. 7 A,B). This may be explained by the observation that peak bases were more prevalent in regions classified as unalignable or structurally variant between the composite and the reference, but non-variant between the SNP-replaced genome and the reference (Supplementary Table 3). Therefore, while fewer peak bases overlapped SNPs and INDELs in the composite WGA than in the SNP-replaced WGA, this advantage was likely outweighed by the greater number of peak bases located in regions that were unalignable or structurally variant relative to the Bowman reference.

**Fig. 7.**
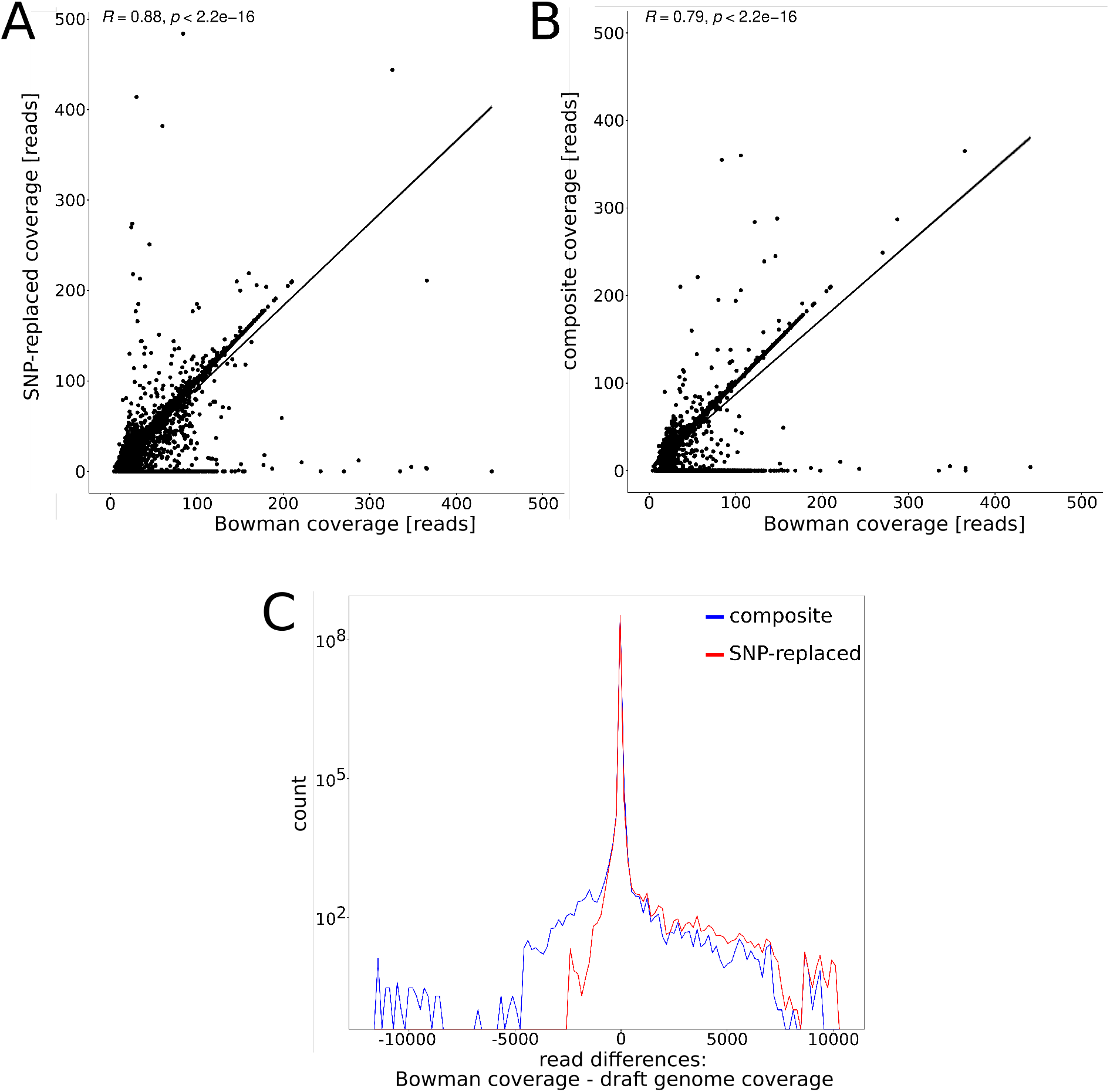
Evaluation of draft genomes based on per base pair comparison of MOA-seq coverage. **A**,**B** Correlation of single-base-pair MOA-seq coverage between each of the draft genomes and the Bowman reference genome. Plotted are only values for bases located within peak summits as called using the Bowman reference sequence. A regression line is plotted in addition and R refers to Spearman’s rho. Axes were limited to a maximum coverage of 500 on each axis. **C** Lines plotted through histograms of the difference in MOA-seq coverage between the Bowman reference sequence and each of the draft genomes at single basepair resolution. The y-axis is logarithmically scaled.

**Table 4.** Correlation of MOA-seq coverage between draft and reference genomes. Spearman’s rho for correlation of per base pair MOA-seq coverage between each genome and the Bowman reference genome. All bases that could be lifted over in at least one genome were considered, with those not lifted over having a coverage of zero. Bases were grouped and subset based on whether they were called as peaks or peak summits in the Bowman reference genome. Analysis was limited to chr1H due to high computational resource requirements otherwise.

| genome | all bases | peak bases | peak summits |
| --- | --- | --- | --- |
| Morex | 0.76 | 0.88 | 0.86 |
| SNP-replaced | 0.82 | 0.90 | 0.88 |
| composite | 0.84 | 0.80 | 0.79 |

The observed differences between the genomes could reflect modest, systematic variation distributed across the genome, or alternatively be driven primarily by a small number of large artefactual differences. To test these hypotheses, we compared the per-bp coverage difference of the MOA-seq alignments directly. Interestingly, the majority of bases, 93 % and 88 % for composite and 89 % and 81 % for the SNP-replaced genome, showed a difference of less than 2 or 1 read, respectively, when compared to the Bowman reference (Fig. 7 C). Despite the overall better performance in this comparison, the composite genome also showed more bases with much higher read coverage (*>*200 reads higher) than the reference (14,732 composite and 10,462 SNP-replaced). Notably, the opposite case, far fewer reads than the reference, was relatively balanced between the composite (5,120 bases) and the SNP-replaced (5,758 bases). Taken together, these results suggest an overall more similar performance of the composite genome to the Bowman reference genome than the SNP-replaced genome, but also an increased number of artefact sites, where the difference in coverage is particularly high.

## Discussion

### Annotation of barley TFBS using MOA-seq

The regulatory space of crop genomes remains poorly characterised. Here, we aimed to provide high-resolution maps of TF footprints of three barley cultivars. We defined the *in vivo cis*-element landscapes of Morex, Bowman and Golden Promise, covering genetic resources, mutant populations, and transformation in barley. The overall number of footprint sites discovered in barley was ~67 % (243k compared to 145k) larger compared to the number of putative *cis*-elements previously found in maize (25). This observation supports a correlation between genome size (5.1 Gbp barley versus 2.1 Gbp maize) and number of potential regulatory regions (46). Additionally, the distribution of barley TF footprints was shifted away from genes compared with maize ((47); see Supplementary Fig. 2).

Comparing the distribution relative to genes, both Self-Transcribing Active Regulatory Region (STARR) sequencing ((48)) and MOA-seq in barley (this study; Fig. 1 A), but also MOA-seq in maize (25), revealed more putative *cis*-elements upstream of genes (TSS +/-50 bp and 5’UTR) than downstream of them (TTS +/-50 bp and 3’UTR).

We note that the 30.2 % average for shared MOA-seq peaks between two cultivars represents a conservative estimate, as peaks were called separately for each genotype, meaning thresholds between signal and noise may differ between genotypes. Therefore, peak calls not shared between genotypes do not always represent statistically significant differences in the occupancy of the genomic region. We observed an overall trend towards a more gene-proximal localisation for these shared peaks (Fig. 2 B), an observation also previously made both *in vitro* in maize (DNA affinity purification sequencing/DAP-seq by Galli et al. (49)) and *in vivo* in barley roots (regions shared between time points; Assay for Transposase-Accessible Chromatin using Sequencing/ATAC-seq by Korinkova et al. (24)).

A plausible explanation of this phenomenon may be the observed decrease in SNPs within shared peaks, as distal regions generally tend to be less conserved than genic or gene-proximal regions and sequence variation can potentially cause differences in TF binding (50). Further investigation into the annotation of the *cis-*regulatory space of barley may allow us to better understand whether our observations are generally applicable to crops with large, complex genomes.

### Comparison of draft genome strategies for comparative TF binding analyses

Annotating regulatory elements of individual cultivars with existing assemblies is feasible (26). Association of sequence variants with TF binding and phenotypic changes, however, generally requires a larger number of genotypes (51, 52), where sequence information is still a limiting factor. We tested both a SNP-replaced and a composite genome strategy as cost-efficient alternatives to classical genome assembly with respect to their ability to analyse TF footprint data e.g., generated by MOA-seq.

For our composite genome strategy, we first verified the PHG’s haplotype calling functionality given selected parameters (Fig. 5). We then quantified the sequence similarity with regard to short variants for both draft genomes compared to the Bowman reference (Fig. 6, Table 2), finding a higher sequence similarity for the composite genome. Larger variants, however, particularly insertions/duplications, were more prevalent in the composite genome compared to the SNP-replaced genome (Table 2). We speculate that these likely originated from sections of selected haplotypes that do not occur in the Bowman reference sequence and were only retained due to being part of the same haplotype (as defined by the PHG) as sections that were supported by Bowman WGS read alignment. This was likely the case for most scaffold haplotypes, but also possible for chromosomal haplotypes.

Furthermore, the composite sequence also contained more bases that could not be aligned to the Bowman reference genome (unaligned bases; Table 2). Unaligned bases unique to the composite-reference WGA occurred mostly in larger sections towards chromosome ends (Supplementary Table 4). In these terminal chromosome regions, varying telomeric repeat content or an increased level of sequence diversity compared to more central regions (27), could have increased the difficulty of accurate WGA during PHG creation. Both reasons could have created inaccuracies in the composite sequence, resulting in failed alignment.

Combining observations of sequence variation and peak-calling accuracy, the most notable difference we found was an increased overlap of annotated MOA-seq peaks with SVs between the composite and reference genome than with SVs between the SNP-replaced and reference genome (Supplementary Table 3). These could be caused by incorrect concatenation of haplotypes or by integration of haplotypes that are only partially supported by short-read alignment. Given that regulatory regions tend to be relatively evolutionarily conserved, this artificial creation of SVs could explain the increase for the composite, while the regulatory regions of the SNP-replaced genome would only be affected by SVs of natural origin and hence less common. This increased overlap with variation in the composite and conservation in the SNP-replaced genome could also explain the observed decrease and increase of the per-base correlation of coverage when limiting the bases to only those annotated as peaks or peak summits for the composite and the SNP-replaced genome, respectively (Table 4).

While the majority of bases showed little difference in coverage, a few displayed extreme differences. Differing from the SNP-replaced genome, the composite contained more bases with extreme coverage relative to a lower reference coverage (Fig. 7 C). These could have been caused by remnants of plastid sequence in the genomes and MOA-seq library or by collapsed repetitive regions accumulating reads, which in turn could either stem from the composite assembly or the genomes of origin for the haplotypes.

Lastly, comparing the ability to call peaks in all three genomes showed that peaks called in draft genomes were broader than in the reference (Table 3). With less accurate genome sequences, some reads were likely not aligned, increasing the difficulty to distinguish between signal and noise and determine the border between them. Additionally, fewer peaks were called using the composite sequence than the other two (Table 3), with the few missing peaks likely stemming from the increased size of unalignable regions between the composite and reference sequence.

Overall, the use of draft genomes facilitated the analysis of MOA-seq data reasonably well given the strong reduction in input data requirement and the resulting observations of strong per-base correlation of coverage, a recovery of called peaks by approximately 80 % and approximately 90 % of the bases between either draft genome and the Bowman reference being non-variant. Analyses using the SNP-replaced genome performed slightly better than using the composite with a higher number of recaptured reference peaks and only a very minimally higher rate of draft-genome-specific peak bases, as well as higher per-base coverage correlation coefficients for bases within peaks. This makes the SNP-replaced genome, together with its simplicity, the preferable option of the two examined strategies to establish draft genomes and could thereby facilitate the analysis of large-scale MOA-seq experiments.

### Potential for improvement of the composite genome assembly strategy

Despite its slightly worse performance compared to the SNP-replaced genome, the composite genome still holds potential, particularly when it comes to the capture of shorter variants or in the case of a more diverse species, where differences between the reference used for the SNP-replaced genome and the genome of interest are larger.

Due to this potential, we want to list potential strategies for improving composite sequence assembly: One option would be to increase the reference-range length from 10 kbp. This could potentially limit the number of SVs created through incorrectly concatenated haplotypes, at the cost of flexibility by limiting haplotype-selection to one genome of origin for longer. A second possibility could be to select the start and end coordinates of the reference ranges more specifically, e.g. to lie within more conserved regions, making the ends of haplotypes more similar to each other and increasing the likelihood of concatenating matching ends. The risk, however, could be the introduction of variants at functionally important regions. A third option could be to build the PHG on genomes assembled using long reads. While the genomes used, assembled using the short-read TRITEX pipeline (37, 53), are already of high quality, assemblies based on long reads could increase contiguity (41), potentially reduce the number of ambiguous “N” bases, and have a better resolution of repetitive regions. A fourth option could be the removal of the concatenated scaffold sequences from the PHG, reducing the number of integrated “N”, but potentially losing important sequences. Alternatively, scaffolds could be added to the graph individually, avoiding the problem of some scaffolds being integrated into the composite, due to the adjacent scaffold being selected by pathfinding. However, this could negatively impact compute performance or interfere with accurate WGA during graph construction. A fifth option to potentially improve the quality and usefulness of the composite sequence could be to increase the number of sequences used to build the PHG, which would increase the variation represented within the graph, at an increased cost. The number of required genomes could potentially be estimated using the saturation curves describing the increase in single-copy pan-genome sequence relative to the number of genomes included in the pan-genome as created by Jayakodi et al. (27). Finally, it would also be possible to not concatenate all of the selected haplotypes, but only to merge those where merging would be supported by overlapping short-reads or paired short-reads. This approach would lead to an overall reduction of the contiguity, but has the potential to improve the accuracy of the sequence, as the risk of introducing variants through incorrect concatenation is mitigated.

In addition to these potential options for improvement, the composite genome assembly strategy can be adapted to preferences and future requirements. In addition to being under active development (54, 55), the PHG offers the option to exchange the used aligner and the variant caller GATK can also be replaced if desired.

Taken together, these points highlight that there are still many avenues to explore to test the potential for further improvement of the composite genome strategy, with respect to both quality and performance.

## Materials and Methods

### Sample preparation and sequencing

Plant material was harvested from *Hordeum vulgare* Morex, Bowman and Golden Promise (three replicates). Plants were grown in a phytochamber at 12 h, 10 °C night and 12 h, 15 °C day. Relative humidity was 60 %, and light at 270 *µ*E (Einstein) during the day. After first tiller bud emergence, the second leaf was harvested 6 h into the day, at 0.5 cm from the leaf base and flash-frozen in liquid nitrogen. Ten plants were pooled per replicate.

MOA-seq and library preparation was performed using 50 mg of homogenised leaf tissue as described in Doan et al. (47) using 37.5 U/mL of MNase. Sequencing was performed using a NovaSeq 6000.

#### Use of publicly available datasets

Assemblies and annotations for Bowman, Golden Promise and Morex (annotation only) were taken from Jayakodi et al. 2024 (GCA_949783105 and GCA_949783185; (27)), and the Morex v3 assembly and annotation from Mascher et al. 2021 (41). Bowman WGS reads were taken from Mayer et al. 2012 (ERR127106, ERR127107, ERR127108, ERR127109, ERR127110, ERR127111, ERR127112, ERR127113; (42)) and those of ten barley cultivars from Weisweiler et al. 2022 (SRR16689792, SRR16689798, SRR16689805, SRR16689797, SRR16689801, SRR16689802, SRR16689796, SRR16689785, SRR16689790, SRR16689783; (44)). The 19 genomes used for the PHG were from Jayakodi et al. 2020 (37).

#### Project-wide use of software tools

Unless otherwise mentioned, R (v4.5.2) was used for analysis (56). Basic file transformations and feature comparisons used AWK and bedtools (57) (v2.30.0). Wgatools maf2chain (v0.1.0; (58)) and CrossMap (v0.7.0; (59)) were used for Multiple Alignment File (MAF) conversion and coordinate translation, respectively (see Supplemental Methods).

#### Analysis of MOA-seq in three barley inbred lines

Sequence preparation and alignment were performed as described by Doan et al. 2026 (47) (see Suppl. Methods). MOA-seq replicate coverage correlation was calculated with deepTools’ (v3.5.4; (60)) multiBamSummary and plotCorrelation with default parameters. MOA-seq coverage was normalised (RPGC) using deepTools bamCoverage (v3.5.4; (60)) and visualised in JBrowse 2 (v4.1.3; (61)) (see also Supplementary Methods).

Peak calling used MACS3 (v3.0.1; (62)) with the three BAM-file replicates merged using samtools as described in Doan et al. 2026 (47), with a q-value cutoff of 0.02. The distance between peaks and the closest gene was determined using bedtools closest. Gene annotation features were extracted using AWK, bedtools sort, merge and subtract using a priority rule in case of multiple feature overlaps and peak-feature overlap determined using bedtools intersect. For all three genomes, the annotation published by Jayakodi et al. 2024 was used (27). For details, see Supplementary Methods.

Genomes were aligned using Anchorwave gff2seq and proali (v1.2.6; (63)) with anchors generated through minimap2 (v2.30-r1287; (64)) using the Morex V3 annotation. Bidirectional translation was achieved through reference and query swapping the MAF files with Martin Firth’s maf-swap script (published with the LAST toolset; (65, 66)). For each genotype and peak, the coordinate of the middle of the peak was then translated to the coordinate system of each of the other two genotypes. A peak was designated as shared between two genotypes, if its middle coordinate translated into the other coordinate system overlapped with a peak called in that genotype and that peak’s middle overlapped with the first peak in its coordinate system. In cases where a peak overlaps with multiple translated peak-middle coordinates or the translated middle of a peak overlaps with multiple peaks, none of the involved peaks were designated as shared.

For the overlap of shared and non-shared peaks between Bowman and Morex with sequence variants and distance to the nearest gene, a window of +/-50 bp around the peak-middle coordinates was used. Variants were extracted from Anchorwave alignments using biokotlin-tools (v0.1.11; (67)) maf-to-gvcf-converter function, and an altered version of Manisha Munashinghe’s R-script (68, 69).

#### Construction of a practical haplotype graph

A practical haplotype graph (PHG; v1.4; (36)) was constructed using mostly default parameters (see PHG Wiki (70)). Barley chromosomes were halved to avoid index size limitations for GATK. Unplaced scaffolds were concatenated with 1000 “N” as spacers to reduce graph complexity. Reference ranges were defined as consecutive 10 kb intervals of the Morex genome.

Pairwise alignments of nineteen barley genomes (37) against Morex were created using the “AssemblyMAFFromAnchorWavePlugin” function (PHG v1.2; Anchorwave v1.0.1). Anchors were generated using a Morex annotation created by Helixer (71, 72) kindly provided by Alisandra Denton. MAFs to GVCF conversion was performed with PHG v1.2, while the remaining steps were performed using v1.4.

#### Pathfinding and generation of draft genome sequences

Adapters of WGS reads of all cultivars (Bowman, CM67; HOR12830; HOR1842; HOR7985; IG128104; IG128216; K10877; Lakhan; Unumli Arpa; W23829/803911, (42, 44)) were trimmed using SeqPurge (ngs-bits v2022_10; (73)) prior to alignment to the graph and path imputation. For details, see Supplementary Methods. After pathfinding, IDs of all haplotypes within the path were extracted using custom Java code, which called the PHG-integrated functions designed for this purpose (bitbucket master-branch accessed on the 13.04.2023; https://bitbucket.org/bucklerlab/practicalhaplotypegraph/src/master/). To validate the determined path, the genome of origin per haplotype was determined and its frequency correlated with the modified Rogers distance of that genome of origin relative to the input WGS reads’ genome (based on a matrix of SNP calls against Morex v3). SNP calls for the genomes of origin were provided by Jayakodi et al. 2020 (37) and the SNP calls of the ten test genotypes were obtained from Weisweiler et al. 2022 (44) (available at: https://doi.org/10.5281/zenodo.6451025) (compare Suppl. Methods).

To create the Bowman preliminary composite sequence, sequences of each haplotype in the path were extracted from their genome of origin, ordered according to corresponding Morex coordinates of each haplotype and concatenated. We then aligned Bowman WGS reads (bwa-mem2 v2.2.1; (74)) to obtain SNPs and INDELs using the Genome Analysis Toolkit (GATK; v4.3.0.0; (43)), which were integrated into the preliminary sequence using bcftools consensus (v1.9; (75)) to finalise the composite genome. For details, see Supplementary Methods.

To create the SNP-replaced genome sequence, variants were called in the same manner, using GATK on Bowman WGS reads aligned against Morex (v3) and SNPs integrated with bcftools consensus. In the same way, variants were also called against the published Bowman sequence (27) as a control.

#### Comparison of draft genome sequences with published reference

SNPs called using WGA between the Bowman and Morex genomes were compared with SNPs called using Bowman WGS against Morex (v3). For each SNP, it was determined whether the composite sequence shared the Bowman genome’s base. WGA was performed using Anchorwave (version 1.0.1; (63); see section “Analysis of MOA-seq in three barley inbred lines”), MAF files were converted to GVCF using the PHG-integrated MAFToGVCFPlugin (PHG version 1.4; (36)) and variants were characterised using Manisha Munashinghe’s script (68, 69). SNPs called from Bowman WGS were taken from the creation of the SNP-replaced genome (see above).

The total sequence of each draft genome was compared to the published Bowman sequence using WGA. WGAs were performed using Anchorwave (v1.2.2; (63)) and variants were called as described above. New variant categories were introduced: “N-SNP” defined as a SNP with the ambiguous base “N” in the query genome and “N-ins” defined as an ambiguous base “N” insertion in the query sequence (length≥1). A comparison of variant annotations by both WGAs was performed by determining the overlap of annotations in the Bowman reference coordinate system. For details, see Supplementary Methods.

#### Comparison of MOA-seq using draft and reference genomes

Bowman MOA-seq reads were aligned to the published Bowman and Morex references and each of the draft genomes (compare “Analysis of MOA-seq in three barley inbred lines”). Peak calling included a control of shortened (60 bp) Bowman WGS reads to alleviate mapping errors. For a per-base comparison of MOA-seq coverage, the Bowman chromosome 1H coordinates were translated to the other genomes, with untranslatable positions counted as zero coverage. Additionally, positions were removed if the difference in coverage between each of the three tested genomes and Bowman was zero.

As a qualitative comparison of the draft genomes’ ability to detect peak regions, the overlap of peaks called using the different genomes was determined using the Bowman reference sequence coordinates. For details, see Supplementary Methods.

## Author declarations

The authors declare that they have no competing interests.

LLM tools were used solely for grammar and language editing; all scientific content was reviewed and approved by the authors.

## Data availability

The data underlying this article are available at the National Center for Biotechnology Information (NCBI) Sequencing Read Archive (SRA) and can be accessed under [tbd]. Draft genomes, MOA-seq coverage and peaks for Bowman, Morex and Golden Promise, as well as the Helixer annotation for Morex and overall code documentation are available at [zenodo; tbd]. The SNP calls provided by the authors of Jayakodi et al. 2020 (37) are available at the European Variation Archive (EVA) under [tbd].

## Author contributions statement

Amelie Kok (Conceptualization [equal], Data curation [equal], Formal analysis [lead], Investigation [lead], Methodology [equal], Software [lead], Validation [lead], Visualization [equal], Writing - original draft [lead], Writing - review & editing [equal]); Jia-Ying Zhu (Investigation [equal], Resources [equal], Writing - review & editing [equal]); Natalie Bajaczyk (Investigation [equal]); Vivien Joisten-Rosenthal (Investigation [supporting], Formal analysis [supporting]); Sohini Mukherjee (Investigation [supporting]); Sergius Weizel (Investigation [supporting]); Venkata Suresh Bonthala (Formal analysis [supporting]); Nafiseh Sargheini (Investigation [supporting]); Bruno Huettel (Methodology [supporting], Writing - review & editing [supporting]); Björn Usadel (Methodology [supporting], Writing - review & editing [supporting]); Julia Engelhorn (Formal analysis [equal], Methodology [equal], Supervision [equal], Validation [equal], Writing - review & editing [equal]); Benjamin Stich (Conceptualization [equal], Funding acquisition [equal], Investigation [equal], Methodology [lead], Software [supporting], Project administration [equal], Resources [equal], Supervision [equal], Validation [equal], Visualization [equal], Writing - original draft [equal], Writing - review & editing [lead]); Thomas Hartwig (Conceptualization [lead], Data curation [equal], Formal analysis [equal], Investigation[lead], Funding acquisition [equal], Methodology [equal], Software [supporting], Project administration [equal], Resources [equal], Supervision [equal], Validation [equal], Visualization [equal], Writing - original draft [equal], Writing - review & editing [lead])

## ACKNOWLEDGEMENTS

We thank Murukarthick Jayakodi (variant calling) and Anne Fiebig (data submission) for providing us with genotype-specific SNP-calls. We thank Alisandra Denton for providing a Helixer annotation for the Morex V3 sequence. We thank Laura Rose and Nathan Springer for their critical feedback and Craig Dent for exploratory discussions. We thank Maximilian Massing for support with Java syntax. We kindly thank the IT service team of the Max Planck Institute for Plant Breeding Research, Cologne, for valuable support. Thank you to Ricardo Henriques for the latex document template.

