## supplemental figures and methods for "A composited genome approach to analyse cistrome diversity in barley"

### 1 Supplementary Figures

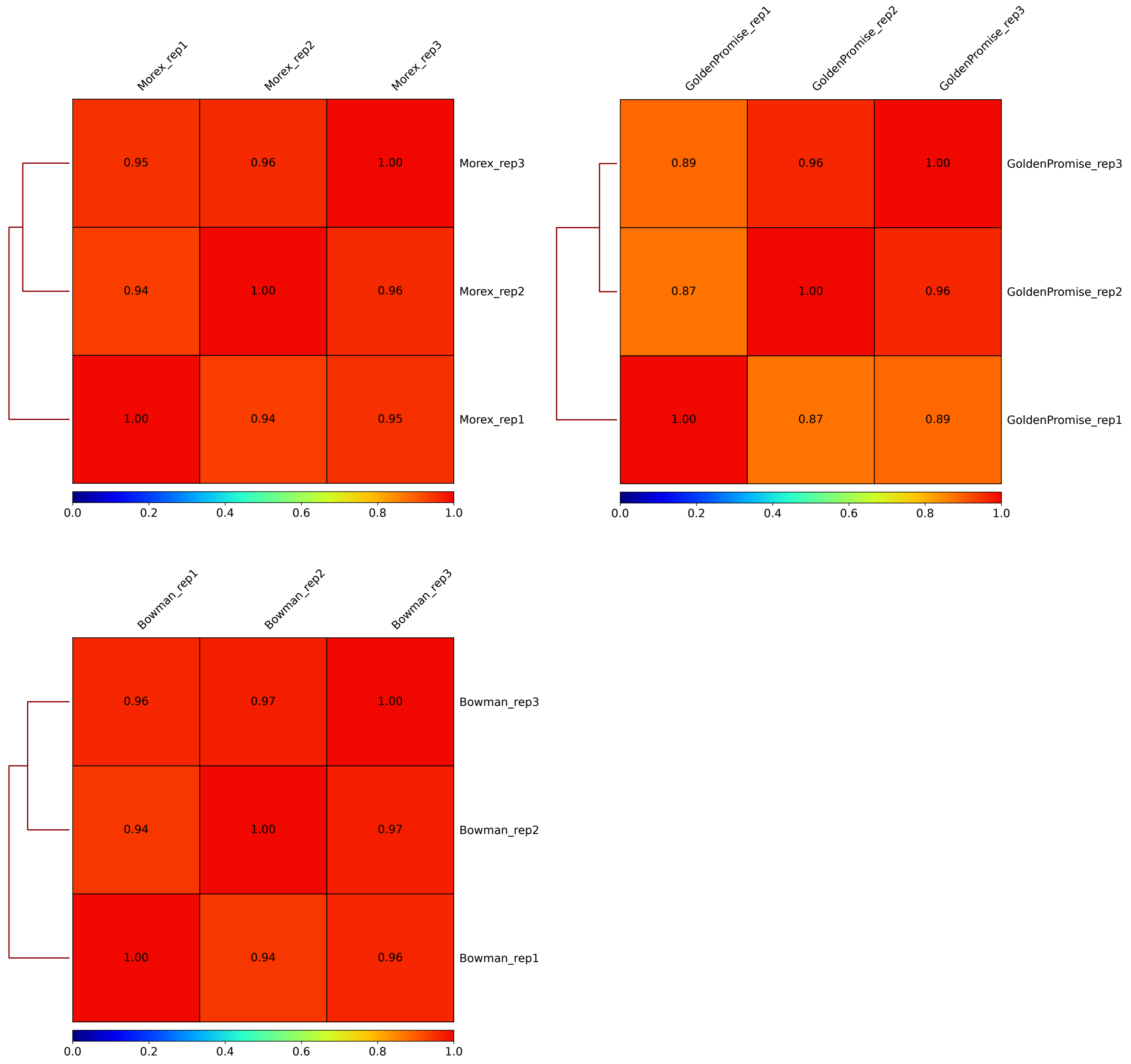


Figure 1: **Correlation of MOA-seq coverage between replicates.** Spearman correlation coefficients for correlations of MOA-seq coverage in 10kb windows between individual replicates of each of the three inbred lines Morex, Bowman and Golden Promise.

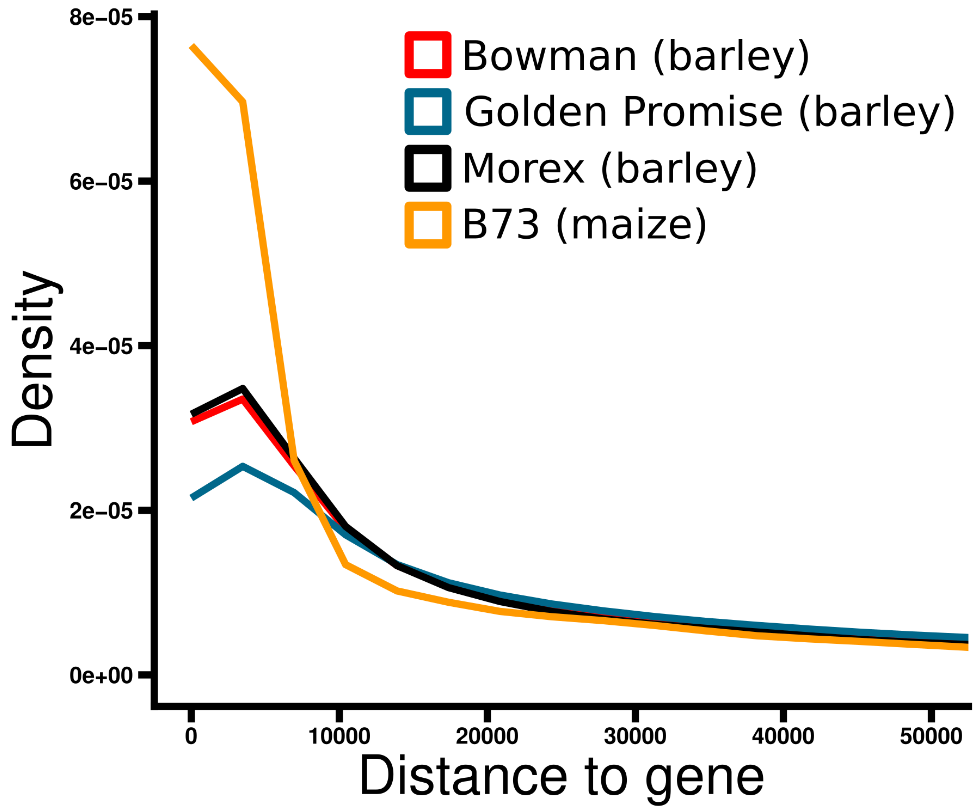


Figure 2: **Localisation of peaks relative to genes.** Relative distance of all peaks to the nearest gene. Excluded are peaks overlapping with genic regions. The x-axis is cut off at 5kb.

#### 2 Supplementary Tables

Table 1: **Counts of peaks categorised by shared-ness between Bowman and Morex and overlap with variants.** Categories of peaks are: "non-shared Bowman" Bowman peaks that are not shared between Morex and Bowman; "non-shared Morex" Morex peaks that are not shared between Morex and Bowman; "shared" peaks that are found to be shared between Bowman and Morex by determining that their middles overlap with the other peak in both coordinate systems; "overl. in Bow. coords." Bowman peaks that are not shared, which still overlap with the middle of a Morex peak in the Bowman coordinate system but whose middle does not overlap with any peak in the Morex coordinate system; "overl. in Mx. coords." Morex peaks that are not shared, which still overlap with the middle of a Bowman peak in the Morex coordinate system but whose middle does not overlap with any peak in the Bowman coordinate system. "overl. too many peaks" if peaks of either genome overlap with too many middles of peaks of the other genotype. These categories are created for all peaks and the subset of peaks that overlap variants between the Bowman and Morex genome. The p-values are of hypergeometric tests for enrichment/depletion (sample: peaks overlapping variants; population: all peaks)

| | all peaks | | peaks overl. variants | | | |
|---|---|---|---|---|---|---|
| **peak category** | **peak number** | **fraction of all crossmapped peaks [%]** | **peak number** | **fraction of all crossmapped peaks [%]** | **p-value depletion** | **p-value enrichment** |
| non-shared Bowman | 136,647 | 37.54 | 36,054 | 39.01 | 1 | 5.45e-27 |
| non-shared Morex | 121,790 | 33.46 | 32,461 | 35.12 | 1 | 1.24e-35 |
| shared | 85,543 | 23.50 | 18,988 | 20.55 | 9.45e-136 | 1 |
| overl. in Bow. coords. | 7,092 | 1.95 | 1,751 | 1.89 | 0.08799 | 0.91202 |
| overl. in Mx. coords. | 7,617 | 2.09 | 1,962 | 2.12 | 0.77740 | 0.22260 |
| overl. too many peaks | 5,323 | 1.46 | 1,201 | 1.30 | 7.38e-07 | 1 |
| sum | 364,012 | 100 | 92,417 | 100 | | |

Table 2: **Alignment statistics for Bowman short-reads against three genomes.** Bases mapped (cigar) refers to the total number of bases that were mapped while filtering based on the CIGAR string so that only matches, inserts, sequence matches and sequence mismatches are counted. Error rate refers to the ratio between mismatches and bases mapped. (See also: samtools stats documentation.

| statistic | Morex | prelim. composite | Bowman |
|---|---|---|---|
| mapped reads | 975,959,779 | 1,031,178,419 | 1,099,947,984 |
| number reads in non-primary alignment | 184,676 | 78,629 | 72,304 |
| bases mapped (cigar) | 97,010,833,339 | 102,578,688,886 | 109,483,567,432 |
| error rate | 8.396653e-03 | 5.455928e-03 | 5.004249e-03 |

Table 3: **Sequence concordance and deviation from the Bowman reference in the two draft genome types.** Bases of the Bowman reference genome were categorised based on which variant-type they were annotated as in the whole genome alignments with the SNP-replaced and the composite genome. In addition to all bases of the genome, those annotated as peaks in the Bowman reference genome are listed in a separate column. Fields' colouration is intended as a visual aid for grouping annotations into similar comparisons.

| **category in composite genome alignment** | **category in SNP-replaced Morex alignment** | **number of bases** | **number of peak bases** |
|---|---|---|---|
| non-variant | non-variant | 3,586,569,860 | 32,152,552 |
| non-variant | SV | 226,108,306 | 1,063,038 |
| SV | non-variant | 187,309,094 | 2,290,999 |
| SV | SV | 113,135,136 | 562,574 |
| non-variant | SNP | 5,261,777 | 30,170 |
| non-variant | INDEL | 3,463,111 | 95,640 |
| SNP | non-variant | 2,467,981 | 24,770 |
| INDEL | non-variant | 1,749,016 | 14,458 |
| non-variant | unaligned | 664,762 | 6,174 |
| unaligned | non-variant | 11,022,326 | 232,847 |
| unaligned | SV | 3,787,884 | 17,342 |
| SV | unaligned | 3,129,083 | 5,580 |
| unaligned | INDEL | 43,987 | 2,243 |
| INDEL | unaligned | 2,815 | 4 |
| unaligned | SNP | 83,808 | 718 |
| SNP | unaligned | 8,404 | 8 |
| SV | SNP | 1,749,207 | 16,129 |
| SV | INDEL | 1,524,337 | 28,773 |
| SNP | SV | 767,152 | 3,198 |
| INDEL | SV | 626,640 | 3,599 |
| SNP | INDEL | 19,667 | 399 |
| INDEL | SNP | 15,810 | 218 |
| INDEL | INDEL | 156,683 | 1,462 |
| SNP | SNP | 194,459 | 1,255 |
| N-SNP | non-variant | 13,199,981 | 146,861 |
| N-SNP | SV | 5,509,332 | 32,550 |
| N-SNP | unaligned | 72,041 | 1,619 |
| N-SNP | SNP | 140,123 | 2,134 |
| N-SNP | INDEL | 90,306 | 2,376 |
| N-SNP | N-SNP | 10,060 | 1,376 |
| non-variant | N-SNP | 53,591 | 1,634 |
| SV | N-SNP | 50,914 | 3,089 |
| unaligned | N-SNP | 80,936 | 0 |
| SNP | N-SNP | 1,304 | 232 |
| INDEL | N-SNP | 646 | 32 |
| N-insertion | non-variant | 5,397,837 | 34,881 |
| N-insertion | SV | 1,698,102 | 2,222 |
| N-insertion | unaligned | 1,143 | 0 |
| N-insertion | SNP | 21,846 | 437 |

Table 4: **Coordinates of regions unalignable between the composite and the Bowman references, but could be aligned between the SNP-replaced genome and the Bowman reference.**

| chromosome | start | stop | length |
|---|---|---|---|
| 1H | 1285609 | 1298612 | 13003 |
| 1H | 472691192 | 472855791 | 164599 |
| 1H | 514507239 | 515719433 | 1212194 |
| 2H | 39567 | 449536 | 409969 |
| 2H | 9472313 | 9611331 | 139018 |
| 2H | 24234576 | 24278299 | 43723 |
| 2H | 550467279 | 550912921 | 445642 |
| 2H | 605345075 | 611510623 | 6165548 |
| 2H | 646714302 | 646769519 | 55217 |
| 3H | 758800 | 1163700 | 404900 |
| 3H | 11496094 | 12052475 | 556381 |
| 3H | 456618549 | 456637534 | 18985 |
| 3H | 586585282 | 587061678 | 476396 |
| 3H | 594414857 | 594708127 | 293270 |
| 3H | 607374537 | 607719829 | 345292 |
| 3H | 623592648 | 624164765 | 572117 |
| 4H | 562606812 | 562801417 | 194605 |
| 4H | 603559292 | 604025300 | 466008 |
| 4H | 606767407 | 607131734 | 364327 |
| 5H | 19882403 | 19973232 | 90829 |
| 5H | 44759783 | 45504083 | 744300 |
| 5H | 500746245 | 500772194 | 25949 |
| 5H | 589522931 | 590003936 | 481005 |
| 6H | 36097728 | 37302611 | 1204883 |
| 6H | 387108865 | 387219459 | 110594 |
| 7H | 604926401 | 604937163 | 10762 |
| 7H | 608205261 | 608214686 | 9425 |

Table 5: **Number of "N" found in selected haplotypes split by chromosome and genome of origin of the haplotype.**

| inbred | chrUn | chr1H | chr2H | chr3H | chr4H | chr5H | chr6H | chr7H |
|---|---|---|---|---|---|---|---|---|
| Akashinriki | 1,728 | 699,098 | 1,675,067 | 1,299,600 | 318,234 | 851,971 | 852,086 | 2,329,007 |
| B1K_04_12 | 0 | 350,739 | 112,401 | 333,659 | 31,451 | 181,189 | 133,796 | 207,563 |
| Barke | 0 | 620,596 | 338,218 | 677,631 | 557,161 | 329,466 | 533,191 | 425,333 |
| Chiba | 4,232 | 42,407 | 180,691 | 71,555 | 11,194 | 80,467 | 33,643 | 144,060 |
| Du li huang | 268,665 | 1,128,144 | 13,204,838 | 1,205,644 | 677,942 | 1,048,884 | 893,237 | 894,891 |
| Golden Promise | 55,375,850 | 242,230 | 606,972 | 240,914 | 292,191 | 205,525 | 221,837 | 148,588 |
| Hockett | 204,292 | 1,510,978 | 774,689 | 820,539 | 3,424,039 | 2,548,434 | 2,620,217 | 3,218,160 |
| HOR10350 | 0 | 305,953 | 267,619 | 250,190 | 290,415 | 258,128 | 333,868 | 449,742 |
| HOR13821 | 0 | 950,402 | 1,195,931 | 1,471,708 | 1,148,997 | 877,683 | 753,988 | 1,282,808 |
| HOR13942 | 7,711 | 521,691 | 692,501 | 939,178 | 514,793 | 862,340 | 720,898 | 2,325,420 |
| HOR21599 | 52,881 | 600,553 | 649,013 | 1,625,985 | 1,111,745 | 2,210,449 | 882,025 | 1,482,269 |
| HOR3081 | 8,075 | 647,618 | 806,137 | 1,394,223 | 492,016 | 727,780 | 679,162 | 1,820,352 |
| HOR3365 | 0 | 1,595,821 | 1,864,569 | 2,983,053 | 1,270,190 | 2,378,365 | 1,459,300 | 4,403,088 |
| HOR7552 | 0 | 234,367 | 646,046 | 357,692 | 149,666 | 474,844 | 322,230 | 955,884 |
| HOR8148 | 1,477,489 | 842,305 | 875,419 | 4,302,594 | 1,550,312 | 5,200,586 | 534,156 | 1,546,344 |
| HOR9043 | 10,713 | 345,764 | 2,233,964 | 1,283,283 | 495,758 | 582,959 | 519,130 | 646,921 |
| Igri | 20,227 | 846,042 | 975,808 | 1,310,255 | 559,097 | 481,344 | 1,575,348 | 931,193 |
| OUN333 | 0 | 393,994 | 2,289,598 | 1,413,453 | 379,396 | 1,198,287 | 363,219 | 1,697,384 |
| RGT Planet | 2 | 2,372,922 | 1,012,643 | 1,372,617 | 1,312,636 | 1,138,023 | 2,656,377 | 646,757 |
| Sum | 57,431,865 | 14,251,624 | 30,402,124 | 23,353,773 | 14,587,233 | 21,636,724 | 16,087,708 | 25,555,764 |

### 3 Supplementary Methods

#### 3.1 Plant material

Plant material was harvested from three different barley (*Hordeum vulgare*) cultivars, namely Morex, Bowman and Golden Promise, in three replicates each. Plants were grown in a phytochamber with cycles of 12 h night and 12 h day, with a nighttime temperature of 10 ℃ and a daytime temperature of 15 ℃. The temperature was gradually increased during the day period. Relative humidity was kept at 60 %, and light intensity at 270 $\mu$E (Einstein) during the day. After the emergence of the first tiller bud, the second leaf was harvested 6 h after the start of the day period, at 0.5 cm distance to the leaf base and immediately flash-frozen with liquid nitrogen. For each replicate the leaves of ten plants were pooled.

#### 3.2 Sample preparation and sequencing

MOA-seq sample and library preparation was performed using 50 mg of frozen and ground leaf tissue following the procedure described in Doan et al. [1]. The MNase concentration was however adjusted to 37.5 U/mL.

##### 3.3 Use of publicly available datasets

For Morex, the version 3 genome sequence and annotation [2] were used for alignment and variant calling. Sequences and annotations for Bowman and Golden Promise were taken from Jayakodi et al. 2024 (GCA_949783105 and GCA_949783185; [3]), as well as another annotation of Morex. Bowman short-read sequencing reads, used to test alternative strategies for genome generation, were taken from Mayer et al. 2012 (ERR127106, ERR127107, ERR127108, ERR127109, ERR127110, ERR127111, ERR127112, ERR127113; [4]). Short-reads of ten further barley cultivars to test the pathfinding algorithm were taken from Weisweiler et al. 2022 (SRR16689792, SRR16689798, SRR16689805, SRR16689797, SRR16689801, SRR16689802, SRR16689796, SRR16689785, SRR16689790, SRR16689783; [5]). In addition to the Morex version 3 sequence, the 19 genomes described by Jayakodi et al. in 2020 were used to build a practical haplotype graph [6].

##### 3.4 Project-wide use of software tools

Unless otherwise mentioned, R version 4.5.2 was used for analysis in R [7]. Packages that were used are ggplot2 (version 4.0.3; [8]), RColorBrewer (version 1.1-3; [9]), dplyr (version 1.2.1; [10]), tidyr (version 1.3.2; [11]), stringr (version 1.6.0 and 1.4.1; [12]), ggpubr (version 0.6.3; [13]), data.table (version 1.18.4; [14]), eulerr (version 7.1.0; [15]). Many basic file transformations and queries were done using custom AWK code. The bedtools suite [16] (v2.30.0), was used to compare features. Translation between coordinate systems, were done using CrossMap (version 0.7.0; [17]) and a Chain file created through MAF-coversion using wgatools maf2chain (version 0.1.0; [18]). Further details for all sections can be found in the Supplemental Methods.

##### 3.5 Analysis of MOA-seq in three barley inbred lines

Sequencing of the MOA-seq libraries of the cultivars Morex, Bowman, and Golden Promise was performed using a NovaSeq 6000 S4 flow cell. Sequencing adaptors were trimmed using SeqPurge (ngs-bits version v2022_10; [19]) with the parameters -min_len 30 -qcut 0. The paired-end reads were then merged into single-end reads using FLASH (version v1.2.11; [20]) with the maximum expected overlap length (-M) set to 100, correcting sequencing errors using base quality scores. Merged reads were then mapped to the corresponding reference genomes using STAR (version 2.7.11a; [21]) with the following parameters: –outSAMmultNmax 1 –outFilterMultimapNmax 4 –winAnchorMultimapNmax 100 –outMultimapperOrder Random –runRNGseed 100 –outBAMsortingBinsN 10 –alignIntronMax 1. Parameters were chosen to limit the number of times a retained read was allowed to map (here: 4), while only retaining a single alignment per read, which is randomly determined in case of draws. alignIntronMax was set to one, to avoid STAR's routines which account for spliced sequences. The resulting bam files were then subset to only contain uniquely mapping reads and reads of up to 80 bp length using samtools (version 1.16.1; [22]) and AWK. Correlations of MOA-seq coverage between the three replicates per genotype were analysed using deeptools' (version 3.5.4; [23]) multiBamSummary function with the "bins" option and default parameters, as well as the plotCorrelation function with the arguments -c spearman -p heatmap –plotNumbers. Visualisations of MOA-seq coverage were created using jbrowse2 (version 4.1.3; [24]). The coverage was extracted from the bam files and normalised to "reads per genomic content" (RPGC) using deeptools bamcoverage (version 3.5.4; [23]) with the following parameters: –exactScaling –smoothLength 0 –binSize 1. The effective genome size for this normalisation was calculated using the unique-kmers.py script of iSeg (versions 1.3.5; [25]) and the average MOA-seq fragment length. The average MOA-seq fragment length was determined using samtools stats with default parameters.

To increase the sensitivity for peak calling, bam files of the three replicates per cultivar were merged using samtools merge. Peak calling was then performed using MACS3 (version v3.0.1; [26]). In order to set the parameters for MACS3, the average MOA-seq fragment length was determined as described above and the genome size was calculated as the total length of sequence in the fasta files. MACS3 parameters were then set as: -q 0.02 -s “AFML” –min-length “AMFL”, –max-gap “AMFL”, –nomodel, –extsize “AMFL”, –keep-dup all, -g “genome size” where AMFL represents the “Average MOA-seq Fragment Length”. The distance between each peak to the closest gene was determined using bedtools (version v2.30.0; [16]) closest, after peaks overlapping with genes were removed using bedtools subtract and custom AWK code. To determine the overlap of peaks with different genic features or intergenic regions, the feature and region coordinates were extracted from the genome annotation using custom AWK code, bedtools sort, merge and subtract. Coordinates were extracted following a set priority list to rule in case of overlaps: 5’UTRs first, then 3’UTRs, exons, introns, 1 kb upstream, 1 kb downstream, 1-5 kb upstream, 1-5 kb downstream and the rest as intergenic. After the coordinates of regions and features were extracted, the overlap was determined using bedtools intersect. For all three genomes the annotation published by Jayakodi et al. 2024 was used [3]. Plotting was performed in R (version 4.5.2; [7]) and ggplot2 (version 4.0.3; [8]).

In order to compare peaks between inbred lines, their genomes were aligned using Anchorwave’s (version 1.2.6; [27]) proali function. For the generation of anchors, the coding sequences were extracted using Anchorwave’s inbuilt gff2seq function and then aligned to the different genomes using minimap2 (version 2.30-r1287; [28]) with Anchorwave’s recommended parameters: -x splice -k 12 -a -p 0.4 -N 20. For the Morex genome, the version 3 of the annotation was used. Anchorwave proali was then run, setting both the reference and query coverage to one (-Q 1 -R 1) and otherwise keeping default parameters. In order to translate coordinates from one genome to another, the resulting MAF files were converted to chain format using wgatools (version 0.1.0; [17]) maf2chain. To create chain files that allow translation in both directions, additional chain files with reference and query genome swapped were created from MAF files converted using Martin Firth’s maf-swap script (published with the LAST toolset; [29, 30]). For each genotype and peak the coordinate of the middle of the peak was then translated to the coordinate system of each of the other two genotypes. This was done using AWK And CrossMap (version 0.7.0; [17]). A peak was designated as shared between two genotypes, if its middle coordinate translated into the other coordinate system overlapped with a peak called in that genotype and that peak's middle overlapped with the first peak in its coordinate system. In cases where a peak overlaps with multiple translated peak-middle coordinates or the translated middle of a peak overlaps with multiple peaks, none of the involved peaks were designated as shared.

To characterise shared and non-shared peaks of the Bowman-Morex comparison further, their overlap with sequence variants and their distance to the nearest genes were determined. A window of +/- 50 bp around the middle coordinates was used to standardise the length of reads. For shared peaks, the middle coordinates of the corresponding peaks were used in each coordinate system. For non-shared peaks the translated coordinates of the middles were used directly. A window of +/- 50 bp around the middle coordinates was then used for the further characterisation of the peaks.
Variants were extracted from the corresponding Anchorwave alignment using biokotlin-tools (version 0.1.11; [31]) maf-to-gvcf-converter function, and altered version of Manisha Munashinghe's R-script [32, 33], as well as custom AWK code. The overlap of peak-windows and variants was determined using bedtools intersect. The distance to the closest genes for the peaks designated as shared and non-shared was taken from the analysis pipeline above. Plots were created in R (version 4.5.2; [7]), using the ggplot2 (version 4.0.3; [8]) and dplyr (version 1.2.1; [10]) packages.

##### 3.6 Construction of a practical haplotype graph

For the creation of a composite genome sequence, a practical haplotype graph (PHG; version 1.4; [34]) was constructed and further used to determine contigs with the highest sequence similarity to 100 bp long Bowman WGS reads (ERR127106, ERR127107, ERR127108, ERR127109, ERR127110, ERR127111, ERR127112, ERR127113; [4]). Default parameters were used when following the pipeline in the PHG Wiki, with a few exceptions described in the following. Due to a limit in chromosome size when using the integrated GATK commands, all barley chromosomes were split in half. In addition, unplaced scaffolds were merged into one contig per assembly by pasting them together with 1000 occurrences of the filler-base N as a separator in order to reduce graph complexity. Instead of splitting reference ranges at the edges of genes, the entire backbone genome (here: Morex v3) was split into ranges of 10kb by specifying the corresponding coordinates in the "validanchors.bed" file. This was done to allow for greater flexibility of the determined path within intergenic regions and increase the likelihood that haplotypes stemming from different assemblies can be picked in order to receive a potentially higher matching sequence to the input WGS reads. The focus on intergenic regions was chosen, due to the biological goal of

this work to identify TFBS. With these changes to the original fasta files and anchor file, the initial database was set up as an SQL database.
In order to fill the graph with haplotypes of diverse barley varieties, 19 genomes assembled by Jayakodi et al. 2020 [6] were aligned to the Morex reference genome in a pairwise fashion. To this end, the “AssemblyMAFFromAnchorWavePlugin” integrated into the PHG (version 1.2) was used. This plugin calls Anchorwave (version 1.0.1) and aligns genomes using default parameters. To generate the anchors, an annotation of the Morex genome created through Helixer and kindly provided by Alisandra Denton was used. After converting the resulting Multiple Alignment Files (MAFs) to Genomic Variant Call Format (GVCF) using the integrated “MAFToGVCFPlugin” (PHG version 1.2), the database was populated with haplotypes of the aligned assemblies using the “LoadHaplotypesFromGVCFPlugin” (PHG version 1.4). The creation of consensus sequences was foregone to capture the most detail in sequence variation.

#### 3.7 Pathfinding and generation of draft genome sequences

The quality of Bowman whole-genome sequencing reads [4], 100 bp in length, were trimmed using SeqPurge (ngs-bits version v2022_10; [19]) setting a minimum length of 30 bp and enabling error correction. All fastq files were then merged into two, one for forward and another for reverse sequencing reads. In the same manner, reads of ten other barley cultivars (CM67; HOR12830; HOR1842; HOR7985; IG128104; IG128216; K10877; Lakhan; Unumli Arpa; W23829/803911, [5]) were trimmed using SeqPurge. This was done to test the pathfinding capability of the PHG. These and the Bowman reads were then aligned to the constructed graph using the ImputePipelinePlugin and imputeTarget pathToVCF option of the PHG software. Most parameters were kept as defaults, with only a few exceptions: The “fParam” parameter was increased to “f15000,16000” to account for the large and repetitive nature of the barley genome. The “minTaxa” parameter, which excludes reference ranges with fewer taxa than the set value, was changed from the default 20 down to one. This was done to retain at least one haplotype sequence for as many reference ranges as possible, so as to not create gaps in the composite sequence where possible. For a similar reason, the “removeEqual” parameter was set to false, so that reference ranges, where ties in the number of reads mapping to two haplotypes occur, are not eliminated. Finally, the minimum transition probability, “minTransitionProb”, was reduced to 0.0005 from the default 0.001, to reduce the number of times where the path transitions from one genotype of origin within the graph to another. This was done to potentially reduce the number of errors that could occur from joining adjacent haplotypes of different origin genotypes.
After the pathfinding step was performed, the IDs of all haplotypes within the path were extracted from the PHG database using custom java code, which in turn called the PHG-integrated functions designed for this purpose (PHG code of the Master-branch of the bitbucket repository was accessed on the 13th of April 2023;
`https://bitbucket.org/bucklerlab/practicalhaplotypegraph/src/master/`). In order

to test the feasibility of the determined path for the ten genotypes mentioned above, the genome of origin of each selected haplotype was determined and set in correlation with the modified Rogers distance of that genome of origin with that of the input WGS reads' genome. The genome of origin for each haplotype was extracted by ID from the "haplotypes" table of the created PHG-sqlite database. The modified Rogers distance was calculated based on a matrix of SNP calls against the Morex genome (v3). The SNP calls of the 19 genomes within the PHG were obtained through communication with the authors of Jayakodi et al. 2020 [6] and the SNP calls of the 10 test-genotypes were obtained through correspondence with the authors of Weisweiler et al. 2022 [5]. The obtained SNP calls in VCF format were merged using bcftools (version 1.9; [35]) isec and merge. The merged list was then filtered to only contain SNPs that were called as homozygous and biallelic within each genotype (no ambiguous SNP calls). Additionally, no more than 8 (of 43) genotypes were allowed to have missing data and the minor allele had to occur for at least 3 of the 43 genotypes. From this filtered list of SNPs, the modified Rogers distance was then computed in R using a custom script and the "stringr" library (R version 4.2.0; stringr version 1.4.1; [12]) and plotted against the relative occurrence of each genome of origin in the determined path (using R (version 4.5.2; [7]), dplyr (version 1.2.1; [10]), ggplot2 (version 4.0.3; [8]), RColorBrewer (version 1.1-3; [9])).

Next a preliminary sequence for the Bowman composite genome was constructed using a custom bash script and bedtools (version v2.30.0; [16]). In brief, the coordinates for each selected haplotype in the path, both in the Morex coordinate system and the coordinate system of the genome of origin, were taken from the PHG sqlite3 database using the extracted haplotype-IDs. Based on the coordinates in the genome of origin, the genomic sequence was extracted using bedtools getfasta (version v2.30.0; [16]) for each haplotype. These sequences were then ordered based on the corresponding Morex coordinates of the haplotype and concatenated into chromosome sequences. To finalise the composite genome sequence SNPs and INDELs were called using the Bowman WGS reads and the preliminary composite sequence and then integrated into the preliminary sequence using bcftools consensus (version 1.9; [35]). The alignment of the reads was performed using bwa-mem2 (version 2.2.1; [36]) and variant calling was done using the Genome Analysis Toolkit (GATK; version version v4.3.0.0; [37]). In more detail: bwa-mem2 was used with the -M option and default parameters. samtools sort and view were used to sort by read name and filter for a minimum mapping quality of 30. samtools fixmate was used to add mate score tags and samtools markdup was used to mark duplicate alignments. The resulting BAM file was fed into GATK HaplotypeCaller and variants were pre-called. The resulting variant calls were then read into the GATK database using GenomicsDBImport and joint genotyping performed using GenotypeGVCFs. On the resulting VCF a first round of hardfiltering was then performed using VariantFiltration with the following filters: "QD < 2.0", "QUAL < 100.0", "SOR > 3.0", "FS > 60.0", "MQ < 40.0", "MQRankSum < -12.5", "ReadPosRankSum < -8.0". Entries which failed to pass these filters were then removed using bcftools view (version 1.11; [35]), while also filtering for homozygous and biallelic variants. After this first round of variant calling with hardfiltering, base recalibration was performed using BaseRecalibrator and ApplyBQSR. A second round of variant calling was then performed using HaplotypeCaller, GenomicsDBImport and GenotypeGVCFs. Variant recalibration based on quality score was done using VariantRecalibrator and ApplyVQSR. The parameters for VariantRecalibrator were "–resource:hapmap3,known=false,training=true,truth=true,prior=15.0", "-tranche 99.5", "-mode BOTH" and the annotations chosen were: "QD", "MQ", "MQRankSum", "ReadPosRankSum", "FS", and "SOR". For ApplyVQSR the truth-sensitivity-filter-level was set to 99.5 to match the truth sensitivity set for VariantRecalibrator.

To create the SNP-replaced genome sequence, variants were called in the same manner, using GATK on an alignment of the Bowman WGS reads against the Morex (v3) genome. Only SNPs were then taken and integrated into the Morex sequence using bcftools consensus.

For an intermediary evaluation step, the number of SNPs and INDELs called against the preliminary composite and the Morex sequence were compared. As a control, variants were also called from an alignment of the Bowman WGS reads against the published Bowman sequence [3]. This was done in the same way as described above.

#### 3.8 Comparison of draft genome sequences with published reference

For a first basic comparison, all SNPs between the published Bowman and Morex sequences were called. To this end, the two genomes were aligned using Anchorwave (version 1.0.1; [27]) using the same parameters as described in section "Analysis of MOA-seq in three barley inbred lines". Variants were called using Manisha Munashinghe's script [32, 33] after the MAF file was converted into GVCF using the MAFToGVCFPlugin tassel plugin integrated into the PHG software (version 1.4; [34]). This set of SNPs was then compared regarding both position and base call with the set of SNPs called using GATK on the Bowman WGS read alignment against the Morex genome. The coordinates of SNPs called using the whole genome alignment were then also translated into the composite genome's coordinate system where the base calls were extracted using bedtools getfasta. These bases were then evaluated based on their match or mismatch with the corresponding base in the published Bowman sequence.

In order to compare the sequences of the SNP-replaced and the composite genome to the published Bowman, both sequences were aligned against the published sequence using the Anchorwave (version 1.2.2; [27]) using the same parameters as described in section "Analysis of MOA-seq in three barley inbred lines". CHAIN files for the translation of coordinates between coordinate systems were created using wgatools (version 0.1.0; [17]). To call variants from the two alignments, the MAF files were converted into GVCF files using the MAFToGVCFPlugin tassel plugin integrated in the PHG (version 1.4; [34]). An altered version of Manisha Munashinghe's script [32, 33] was then used to extract and characterise variants. In particular, the category of "N-SNP" and "N-ins" were added. We defined "N-SNP" as a SNP that had a base change that resulted in a call of the non-descript base "N" in the query genome. "N-ins" was defined as an insertion in the query sequence consisting only of the non-descript base "N". The number of variant bases of each type were determined using custom AWK code. A comparison of variant annotations for every base in the published Bowman genome sequence was done by converting the output of Manisha Munashinghe's script into bed files using the published Bowman coordinate system and using bedtools intersect to determine which annotations overlapped. A summary of this result was then determined using custom AWK and R code (version 4.5.2 [7]; dplyr version 1.2.1 [10]; tidyr version 1.3.2 [11]).

For regions missing in the composite genome, but present in the SNP-replaced genome, we tested if the reason for their absence in the composite sequence was due to no haplotype being selected for the corresponding reference range. To this end, we took the coordinates of the missing regions from the SNP-replaced genome and compared them to the merged coordinates of missing reference ranges in the Bowman path through the PHG (AWK and bedtools used; Note that the reference ranges are in the Morex coordinate system and the SNP-replaced genome shares its coordinate system with Morex).
The origin of the non-descript "N" bases in the composite sequence was determined by counting the number of "N" for every extracted sequence of a haplotype (compare section "Pathfinding and generation of draft genome sequences"), grouped by genome of origin and chromosome. In addition, the number of "N" was also counted in the finalised composite sequence to test how many "N" were removed through the variant replacement step.

#### 3.9 Comparison of MOA-seq using draft and reference genomes

Bowman MOA-seq reads were aligned to the published Bowman reference sequence and each of the two draft genomes sequences. The resulting coverage tracks were then evaluated regarding the similarity of the tracks of the draft genomes to the track of the published Bowman reference sequence. As an additional baseline/control, the Bowman MOA-seq reads were also aligned against the Morex reference sequence. The read alignment to the draft genomes was performed using STAR as described above (see section "Analysis of MOA-seq in three barley inbred lines"), with the only exception being that only unique mapping reads were retained by setting the "outFilterMultimapNmax" parameter to one. In the method described above, this was instead done using a filter step after the STAR read alignment. The MOA-seq coverage was determined using deeptools bamCoverage (version 3.5.4; cite). The MOA-seq coverage per single base pair of chromosome 1H was then compared between the Bowman reference, and the two draft genomes and the Morex genome. To this end, every base of the Bowman reference sequence of chromosome 1H was translated into the coordinate system of each of the other genomes, where possible. In cases where no coverage value could be determined for one genome (i.e. if that position could not be translated between coordinate systems), the value was set to zero. In addition, every position was removed where the difference in coverage between each of the three tested genomes and the Bowman reference sequence was zero. The Spearman correlation coefficient for the correlation of the coverage in the Bowman reference sequence and each of the three tested genomes was calculated in R (version 4.5.2 [7]; dplyr version 1.2.1 [10]; tidyr version 1.3.2 [11]).

To focus particularly on MOA-seq peak-regions, peaks were called to above, using the alignment of MOA-seq reads against the Bowman reference sequence and the two draft genomes. Peaks were called using MACS3, using the same parameters as described in section "Analysis of MOA-seq in three barley inbred lines", but with the addition of a control (-c). This control was the alignment of Bowman WGS reads against each of the genomes in the same manner as the MOA-seq reads, after the WGS reads had been shortened to the average length of MOA-seq fragments of 60 bp. A control was included to minimise the number of false peak calls caused by sequence artifacts. The correlation of coverage at single-basepair resolution, as described above, was repeated once for bases located in peaks called based on the Bowman reference sequence alignment, and another time for bases located in those peaks' summits.

As a qualitative comparison of the draft genomes' ability to detect peak regions, the overlap of peaks called using the different genomes was determined. For this purpose, the coordinates of the peaks called using the composite genome and of the peaks called using the SNP-replaced genome were translated to the Bowman reference genome coordinate system using CrossMap (version 0.7.0; [17]). The number of peak-bases of each of the draft genomes that overlapped with peak-bases of the Bowman reference sequence was determined using bedtools intersect. The number of true positive peak-bases was then calculated as the number of bases overlapping between the draft genome peaks and the Bowman reference peaks, while the number of false-positive bases was calculated as the number of peak-bases called in each draft genome that was not called using the Bowman reference sequence.

[2] Martin Mascher, Thomas Wicker, Jerry Jenkins, Christopher Plott, Thomas Lux, Chu Shin Koh, Jennifer Ens, Heidrun Gundlach, Lori B Boston, Zuzana Tulpová, Samuel Holden, Inmaculada Hernández-Pinzón, Uwe Scholz, Klaus F X Mayer, Manuel Span-

nagl, Curtis J Pozniak, Andrew G Sharpe, Hana Šimková, Matthew J Moscou, Jane Grimwood, Jeremy Schmutz, and Nils Stein. “Long-read sequence assembly: a technical evaluation in barley”. In: *The Plant Cell* 33.6 (Mar. 2021), pp. 1888–1906.

[3] Murukarthick Jayakodi, Qiongxian Lu, Hélène Pidon, M. Timothy Rabanus-Wallace, Micha Bayer, Thomas Lux, Yu Guo, Benjamin Jaegle, Ana Badea, Wubishet Bekele, Gurcharn S. Brar, Katarzyna Braune, Boyke Bunk, Kenneth J. Chalmers, Brett Chapman, Morten Egevang Jørgensen, Jia-Wu Feng, Manuel Feser, Anne Fiebig, Heidrun Gundlach, Wenbin Guo, Georg Haberer, Mats Hansson, Axel Himmelbach, Iris Hoffie, Robert E. Hoffie, Haifei Hu, Sachiko Isobe, Patrick König, Sandip M. Kale, Nadia Kamal, Gabriel Keeble-Gagnère, Beat Keller, Manuela Knauft, Ravi Koppolu, Simon G. Krattinger, Jochen Kumlehn, Peter Langridge, Chengdao Li, Marina P. Marone, Andreas Maurer, Klaus F. X. Mayer, Michael Melzer, Gary J. Muehlbauer, Emiko Murozuka, Sudharsan Padmarasu, Dragan Perovic, Klaus Pillen, Pierre A. Pin, Curtis J. Pozniak, Luke Ramsay, Pai Rosager Pedas, Twan Rutten, Shun Sakuma, Kazuhiro Sato, Danuta Schüler, Thomas Schmutzer, Uwe Scholz, Miriam Schreiber, Kenta Shirasawa, Craig Simpson, Birgitte Skadhauge, Manuel Spannagl, Brian J. Steffenson, Hanne C. Thomsen, Josquin F. Tibbits, Martin Toft Simmelsgaard Nielsen, Corinna Trautewig, Dominique Vequaud, Cynthia Voss, Penghao Wang, Robbie Waugh, Sharon Westcott, Magnus Wohlfahrt Rasmussen, Runxuan Zhang, Xiao-Qi Zhang, Thomas Wicker, Christoph Dockter, Martin Mascher, and Nils Stein. “Structural variation in the pangenome of wild and domesticated barley”. In: *Nature* 636.8043 (Dec. 2024), pp. 654–662.

[4] Klaus F. X. Mayer, Robbie Waugh, Peter Langridge, Timothy J. Close, Roger P. Wise, Andreas Graner, Takashi Matsumoto, Kazuhiro Sato, Alan Schulman, Gary J. Muehlbauer, Nils Stein, Ruvini Ariyadasa, Daniela Schulte, Naser Poursarebani, Ruonan Zhou, Burkhard Steuernagel, Martin Mascher, Uwe Scholz, Bujun Shi, Kavitha Madishetty, Jan T. Svensson, Prasanna Bhat, Matthew Moscou, Josh Resnik, Pete Hedley, Hui Liu, Jenny Morris, Zeev Frenkel, Avraham Korol, Hélène Bergès, Stefan Taudien, Marius Felder, Marco Groth, Matthias Platzer, Axel Himmelbach, Stefano Lonardi, Denisa Duma, Matthew Alpert, Francesa Cordero, Marco Beccuti, Gianfranco Ciardo, Yaqin Ma, Steve Wanamaker, Federica Cattonaro, Vera Vendramin, Simone Scalabrin, Slobodanka Radovic, Rod Wing, Michele Morgante, Thomas Nussbaumer, Heidrun Gundlach, Mihaela Martis, Jesse Poland, Manuel Spannagl, Matthias Pfeifer, Cédric Moisy, Jaakko Tanskanen, Andrea Zuccolo, Joanne Russell, Arnis Druka, David Marshall, Micha Bayer, David Swarbreck, Dharanya Sampath, Sarah Ayling, Melanie Febrer, Mario Caccamo, Tsuyoshi Tanaka, Steve Wannamaker, Thomas Schmutzer, John W. S. Brown, Geoffrey B. Fincher, The International Barley Genome Sequencing Consortium, Principal investigators, Physical map construction, direct anchoring direct, Genomic sequencing, assembly, BAC sequencing, BAC-end sequencing, Integration of physical/genetic map, sequence resources sequence, Gene annotation, Repetitive DNA analysis, Transcriptome sequencing, analysis, Re-sequencing, diversity analysis diversity, Writing, and editing of the manuscript editing of the. “A physical,

genetic and functional sequence assembly of the barley genome". In: *Nature* 491.7426 (Nov. 2012), pp. 711–716.

[5] Marius Weisweiler, Christopher Arlt, Po-Ya Wu, Delphine Van Inghelandt, Thomas Hartwig, and Benjamin Stich. "Structural variants in the barley gene pool: precision and sensitivity to detect them using short-read sequencing and their association with gene expression and phenotypic variation". en. In: *Theor. Appl. Genet.* 135.10 (Oct. 2022), pp. 3511–3529.

[6] Murukarthick Jayakodi, Sudharsan Padmarasu, Georg Haberer, Venkata Suresh Bonthala, Heidrun Gundlach, Cécile Monat, Thomas Lux, Nadia Kamal, Daniel Lang, Axel Himmelbach, Jennifer Ens, Xiao-Qi Zhang, Tefera T. Angessa, Gaofeng Zhou, Cong Tan, Camilla Hill, Penghao Wang, Miriam Schreiber, Lori B. Boston, Christopher Plott, Jerry Jenkins, Yu Guo, Anne Fiebig, Hikmet Budak, Dongdong Xu, Jing Zhang, Chunchao Wang, Jane Grimwood, Jeremy Schmutz, Ganggang Guo, Guoping Zhang, Keiichi Mochida, Takashi Hirayama, Kazuhiro Sato, Kenneth J. Chalmers, Peter Langridge, Robbie Waugh, Curtis J. Pozniak, Uwe Scholz, Klaus F. X. Mayer, Manuel Spannagl, Chengdao Li, Martin Mascher, and Nils Stein. "The barley pangenome reveals the hidden legacy of mutation breeding". In: *Nature* 588.7837 (Dec. 2020), pp. 284–289.

[7] R Core Team. *R: A Language and Environment for Statistical Computing*. R Foundation for Statistical Computing. Vienna, Austria, 2025.

[8] Hadley Wickham. *ggplot2: Elegant Graphics for Data Analysis*. Springer-Verlag New York, 2016.

[9] Erich Neuwirth. *RColorBrewer: ColorBrewer Palettes*. R package version 1.1-3. 2022.

[10] Hadley Wickham, Romain François, Lionel Henry, Kirill Müller, and Davis Vaughan. *dplyr: A Grammar of Data Manipulation*. R package version 1.2.1. 2026.

[11] Hadley Wickham, Davis Vaughan, and Maximilian Girlich. *tidyr: Tidy Messy Data*. R package version 1.3.2. 2025.

[12] Hadley Wickham. *stringr: Simple, Consistent Wrappers for Common String Operations*. R package version 1.6.0. 2025.

[13] Alboukadel Kassambara. *ggpubr: 'ggplot2' Based Publication Ready Plots*. R package version 0.6.3. 2026.

[14] Tyson Barrett, Matt Dowle, Arun Srinivasan, Jan Gorecki, Michael Chirico, Toby Hocking, Benjamin Schwendinger, and Ivan Krylov. *data.table: Extension of 'data.frame'*. R package version 1.18.4. 2026.

[15] Johan Larsson and Peter Gustafsson. "A Case Study in Fitting Area-Proportional Euler Diagrams with Ellipses Using eulerr". In: *Proceedings of International Workshop on Set Visualization and Reasoning*. Vol. 2116. Edinburgh, United Kingdom: CEUR Workshop Proceedings, Apr. 2018, pp. 84–91.

[16] Aaron R. Quinlan and Ira M. Hall. "BEDTools: a flexible suite of utilities for comparing genomic features". In: *Bioinformatics* 26.6 (Mar. 2010), pp. 841–842.

[17] Hao Zhao, Zhifu Sun, Jing Wang, Haojie Huang, Jean-Pierre Kocher, and Liguo Wang. "CrossMap: a versatile tool for coordinate conversion between genome assemblies". en. In: *Bioinformatics* 30.7 (Apr. 2014), pp. 1006–1007.

[18] Wenjie Wei, Songtao Gui, Jian Yang, Erik Garrison, Jianbing Yan, and Hai-Jun Liu. "wgatools: an ultrafast toolkit for manipulating whole-genome alignments". In: *Bioinformatics* 41.4 (Apr. 2025), btaf132.

[19] Marc Sturm, Christopher Schroeder, and Peter Bauer. "SeqPurge: highly-sensitive adapter trimming for paired-end NGS data". en. In: *BMC Bioinformatics* 17.1 (May 2016), p. 208.

[20] Tanja Magoč and Steven L. Salzberg. "FLASH: fast length adjustment of short reads to improve genome assemblies". In: *Bioinformatics* 27.21 (Nov. 2011), pp. 2957–2963.

[21] Alexander Dobin, Carrie A Davis, Felix Schlesinger, Jorg Drenkow, Chris Zaleski, Sonali Jha, Philippe Batut, Mark Chaisson, and Thomas R Gingeras. "STAR: ultrafast universal RNA-seq aligner". en. In: *Bioinformatics* 29.1 (Jan. 2013), pp. 15–21.

[22] Heng Li, Bob Handsaker, Alec Wysoker, Tim Fennell, Jue Ruan, Nils Homer, Gabor Marth, Goncalo Abecasis, Richard Durbin, and 1000 Genome Project Data Processing Subgroup. "The Sequence Alignment/Map format and SAMtools". In: *Bioinformatics* 25.16 (Aug. 2009), pp. 2078–2079.

[23] Fidel Ramírez, Devon P Ryan, Björn Grüning, Vivek Bhardwaj, Fabian Kilpert, Andreas S Richter, Steffen Heyne, Friederike Dündar, and Thomas Manke. "deepTools2: a next generation web server for deep-sequencing data analysis". In: *Nucleic Acids Research* 44.W1 (July 2016), W160–W165.

[24] Colin Diesh, Garrett J. Stevens, Peter Xie, Teresa De Jesus Martinez, Elliot A. Hershberg, Angel Leung, Emma Guo, Shihab Dider, Junjun Zhang, Caroline Bridge, Gregory Hogue, Andrew Duncan, Matthew Morgan, Tia Flores, Benjamin N. Bimber, Robin Haw, Scott Cain, Robert M. Buels, Lincoln D. Stein, and Ian H. Holmes. "JBrowse 2: a modular genome browser with views of synteny and structural variation". In: *Genome Biology* 24.1 (Apr. 2023), p. 74.

[25] Senthil B. Girimurugan, Yuhang Liu, Pei-Yau Lung, Daniel L. Vera, Jonathan H. Dennis, Hank W. Bass, and Jinfeng Zhang. "iSeg: an efficient algorithm for segmentation of genomic and epigenomic data". In: *BMC Bioinformatics* 19.1 (Apr. 2018), p. 131.

[26] Yong Zhang, Tao Liu, Clifford A. Meyer, Jérôme Eeckhoute, David S. Johnson, Bradley E. Bernstein, Chad Nusbaum, Richard M. Myers, Myles Brown, Wei Li, and X. Shirley Liu. "Model-based Analysis of ChIP-Seq (MACS)". In: *Genome Biology* 9.9 (Sept. 2008), R137.

[35] Petr Danecek, James K Bonfield, Jennifer Liddle, John Marshall, Valeriu Ohan, Martin O Pollard, Andrew Whitwham, Thomas Keane, Shane A McCarthy, Robert M Davies, and Heng Li. “Twelve years of SAMtools and BCFtools”. In: *GigaScience* 10.2 (Feb. 2021). giab008.

[36] Md. Vasimuddin, Sanchit Misra, Heng Li, and Srinivas Aluru. “Efficient Architecture-Aware Acceleration of BWA-MEM for Multicore Systems”. In: *2019 IEEE International Parallel and Distributed Processing Symposium (IPDPS)*. 2019, pp. 314–324.

[37] Aaron McKenna, Matthew Hanna, Eric Banks, Andrey Sivachenko, Kristian Cibulskis, Andrew Kernytsky, Kiran Garimella, David Altshuler, Stacey Gabriel, Mark Daly, and Mark A. DePristo. “The Genome Analysis Toolkit: A MapReduce framework for analyzing next-generation DNA sequencing data”. In: *Genome Research* (2010).
